# Effects-sizes of deletions and duplications on autism risk across the genome

**DOI:** 10.1101/2020.03.09.979815

**Authors:** Elise Douard, Abderrahim Zeribi, Catherine Schramm, Petra Tamer, Mor Absa Loum, Sabrina Nowak, Zohra Saci, Marie-Pier Lord, Borja Rodríguez-Herreros, Martineau Jean-Louis, Clara Moreau, Eva Loth, Gunter Schumann, Zdenka Pausova, Mayada Elsabbagh, Laura Almasy, David C. Glahn, Thomas Bourgeron, Aurélie Labbe, Tomas Paus, Laurent Mottron, Célia M. T. Greenwood, Guillaume Huguet, Sébastien Jacquemont

## Abstract

**Objective:** Deleterious copy number variants (CNVs) are identified in up to 20% of individuals with autism. However, only 13 genomic loci have been formally associated with autism because the majority of CNVs are too rare to perform individual association studies. To investigate the implication of undocumented CNVs in neurodevelopmental disorders, we recently developed a new framework to estimate their effect-size on intelligence quotient (IQ) and sought to extend this approach to autism susceptibility and multiple cognitive domains.

**Methods:** We identified CNVs in two autism samples (Simons Simplex Collection and MSSNG) and two unselected populations (IMAGEN and Saguenay Youth Study). Statistical models integrating scores of genes encompassed in CNVs were used to explain their effect on autism susceptibility and multiple cognitive domains.

**Results:** Among 9 scores of genes, the “probability-of-being loss-of-function intolerant” (pLI) best explains the effect of CNVs on IQ and autism risk. Deletions decrease IQ by a mean of 2.6 points per point of pLI. The effect of duplications on IQ is three-fold smaller. The odd ratios for autism increases when deleting or duplicating any point of pLI. This increased autism risk is similar in subgroups of individuals below or above median IQ. Once CNV effects on IQ are accounted for, autism susceptibility remains mostly unchanged for duplications but decreases for deletions. Model estimates for autism risk overlap with previously published observations. Deletions and duplications differentially affect social communication, behaviour, and phonological memory, whereas both equally affect motor skills.

**Conclusions:** Autism risk conferred by duplications is less influenced by IQ compared to deletions. CNVs increase autism risk similarly in individuals with high and low IQ. Our model, trained on CNVs encompassing >4,500 genes, suggests highly polygenic properties of gene dosage with respect to autism risk. These models will help interpreting CNVs identified in the clinic.

## INTRODUCTION

Autism is a neurodevelopmental condition currently defined by atypical social communication and interaction, intense interests, and repetitive behaviour (1). Levels of general intelligence and language are not diagnostic criteria but are recognized as clinical specifiers which have been defined as important features of the heterogeneity of autism. (2) Neurodevelopmental and psychiatric comorbidities occur in up to 70% of children with autism. (3) The heritability of autism has been estimated between 50-80%. (4, 5) Deleterious single nucleotide variants (SNV) and copy number variants (CNVs) are identified in 15 to 20% of individuals with autism. (6–8) The largest rare variant autism case-control association studies to date have formally associated 91 genes and CNVs at only 13 genomic loci. (9–13) Many more genomic loci are likely implicated as suggested by the overall increase in CNV burden associated with autism. (9, 12, 14–16) Therefore, the susceptibility to autism conferred by most CNVs remains undocumented. This is particularly problematic in the neurodevelopmental clinic, where undocumented CNVs are routinely diagnosed in a large proportion of patients.

Even less is known about the effect-size of CNVs on the cognitive and behavioural dimensions related to autism, which have only been characterized for a handful of recurrent CNVs (*e.g.* 22q11.2, 16p11.2, 15q11.2, and 1q21.1 loci). These CNVs show reproducible effect-sizes on cognition, language, socio-communication, and brain structure, suggesting that these alterations drive their over-representation in autism or other neurodevelopmental and psychiatric conditions. (17–19)

Limited progress has been made in identifying phenotype-genotype relationships in autism. Studies have demonstrated that rare *de-novo* variants are associated with lower intelligence quotient (IQ) and are over-represented in females. (16, 20–23) *De novo* variants have also been associated with an atypical autism profile characterized by less impairment in social communication and language, as well as greater likelihood of motor delay. (24, 25) Overall, the reasons underlying the overrepresentation of rare variants in autistic individuals remains unclear. It may be due to their effect on core symptoms of autism, or DSM-5-defined clinical specifiers of autism (intelligence, language, co-occurring conditions). Since CNVs studied to date have not been specifically associated with autism, it would be expected that cognitive and behavioral dimensions affected by CNVs are relevant for other neurodevelopmental and psychiatric disorders.

We previously reported that statistical models, trained on benign deletions in populations not selected for a clinical condition, can accurately estimate the effect-size of deleterious deletions on non-verbal IQ (NVIQ). (26) These results suggest that 1) the effect-size of deletions on NVIQ can be estimated using constraint scores, such as the “probability of being Loss-of-function Intolerant” (pLI, definition in textbox, Figure 1) (27), and 2) the effect of haploinsufficiency on NVIQ applies to a large proportion of the genome, consistent with a highly polygenic model. (28, 29) Using pLI as an explanatory variable, we estimated that one third of the coding genes affect NVIQ by >1 point, when deleted. (26) Previously, we were unable to establish the effect-size of duplications, likely due to inadequate power with the then-available sample size. Here, we propose to develop similar models to estimate autism susceptibility conferred by undocumented CNVs. We also aim to estimate their effects on cognitive and behavioural dimensions, which may underpin their overrepresentation in autism.

**Figure 1.**
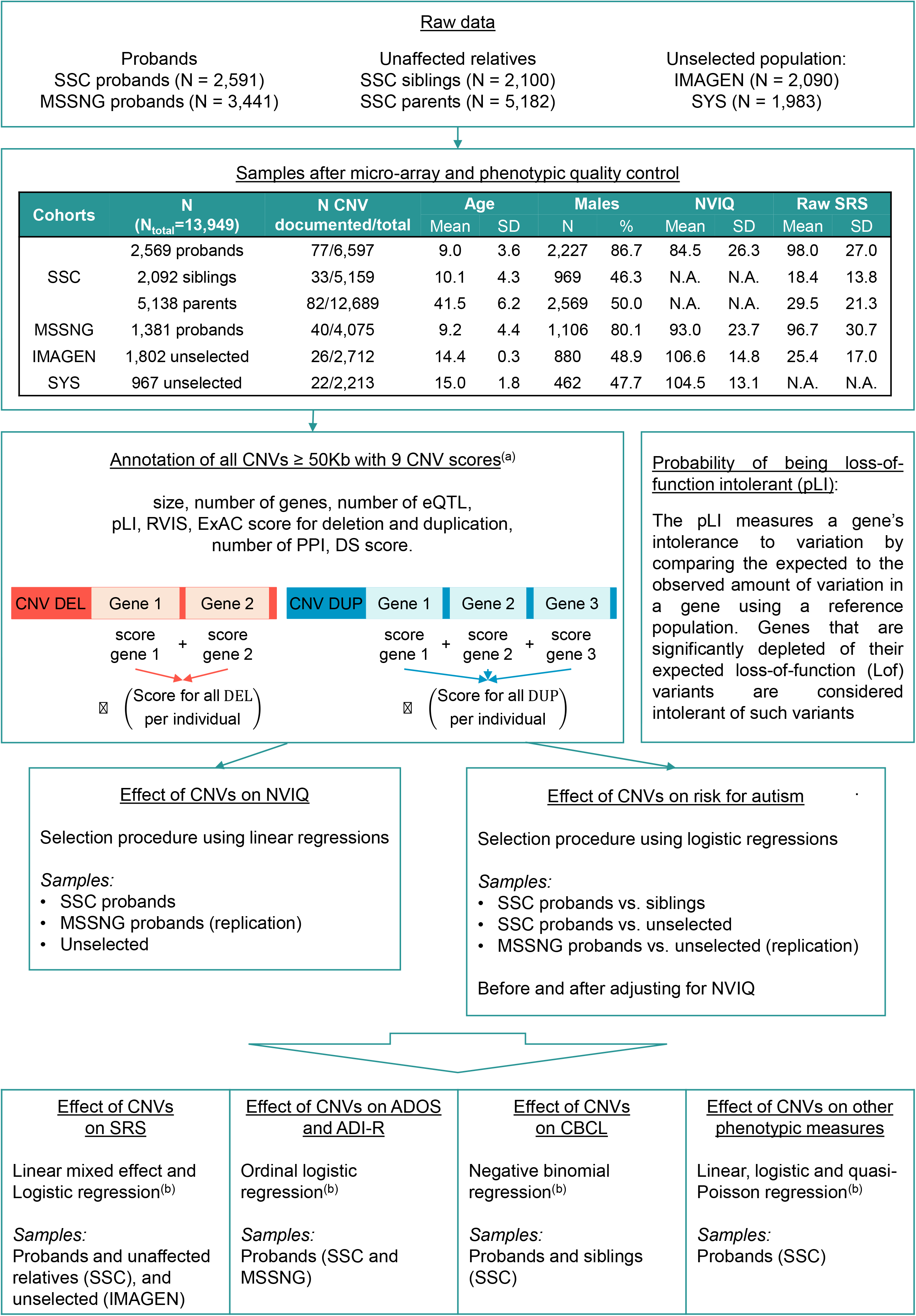
Methodological pipeline. Probands from the SSC and MSSNG are defined as individuals recruited on the basis of a diagnosis of autism. Siblings and parents from the SSC did not meet diagnostic criteria for autism. 10 individuals from IMAGEN and none in SYS met criteria for ASD (as estimated by the Development and Well-Being Assessment, DAWBA). 1,490 unaffected siblings from the SSC, 3,660 unaffected parents from the SSC and 1,465 individuals from the general population carry at least one CNV ≥ 50kb (Table S1 in the online supplement). (a) Microarray quality control and CNV selection and annotation were performed as previously published (15) (Methods in the online supplement); (b) The model used and available data for each phenotype are detailed in Table 1 and Tables S7, S8, S9, and S10 in the online supplement. SSC: Simon Simplex Collection; SYS: Saguenay Youth Study; CNV: copy number variants; SD: standard deviation; N.A.: Not applicable; NVIQ: non-verbal intelligence quotient; SRS: Social Responsiveness Scale; ADOS: Autism Diagnostic Observation Schedule; ADI-R: Autism Diagnostic Interview-Revised; CBCL: Child behaviour Checklist.

We 1) tested whether the effect-size of gene dosage on NVIQ is the same across unselected populations and autism cohorts, 2) selected models that best explain the autism risk conferred by any deletions or duplications, while accurately adjusting for their effect on NVIQ established in step 1, and 3) investigated the cognitive, behavioural, and motor phenotypes that may explain the association between gene dosage and autism.

**Table 1:**
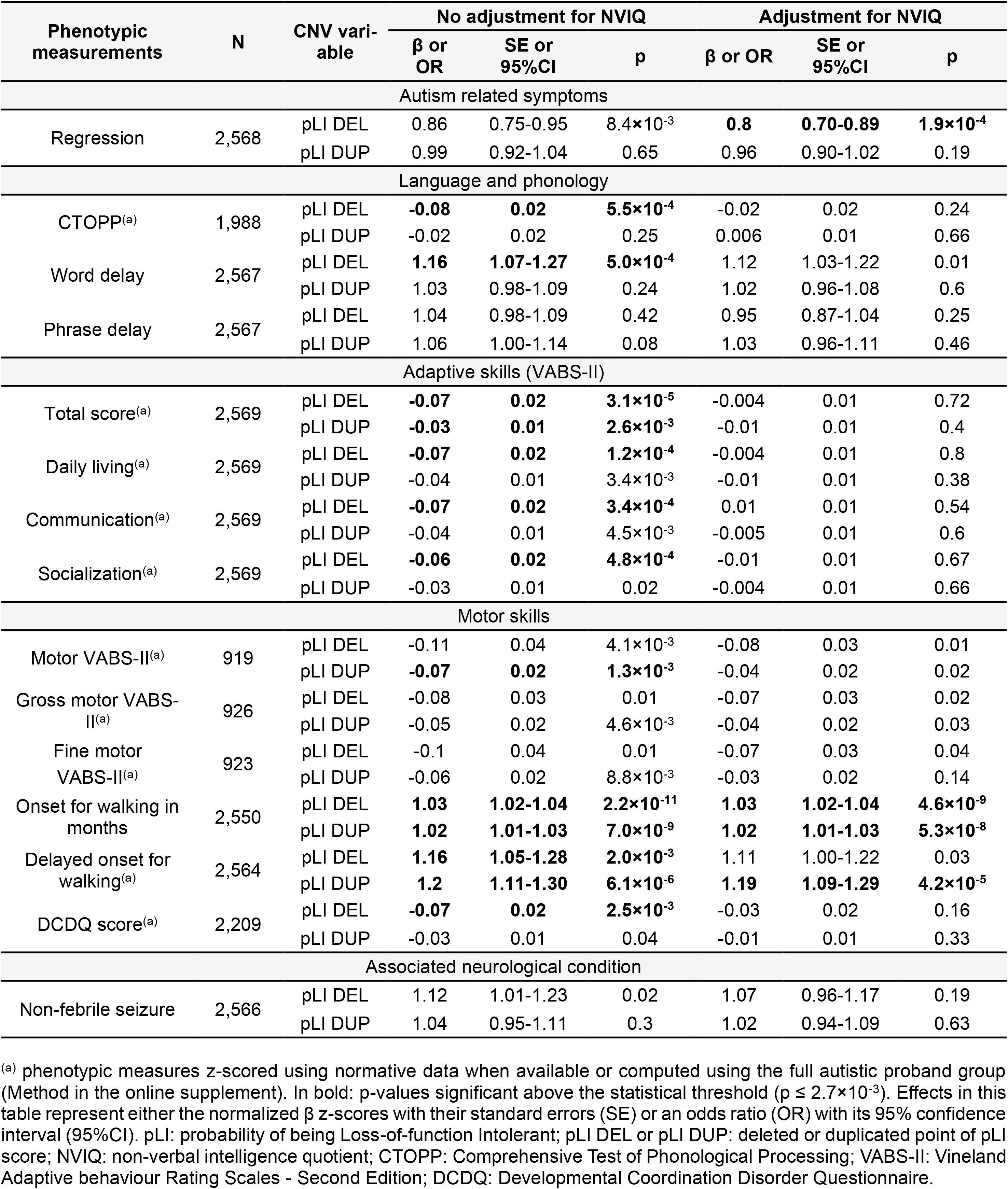
Effect-size of gene dosage measured by pLI on phenotypes in autistic probands from the SSC.

Models integrating genomic and functional scores of genes included in CNVs were trained on all CNVs ≥ 50 kb identified in two autism cohorts and two cohorts recruited from unselected populations. We provide a novel framework to model autism risk and the phenotypic profile of rare variants, regardless of effect-size and inheritance. This approach contrasts with previous genotype-phenotype studies restricted to small groups of individuals with *de-novo* or recurrent variants.

## METHODS

### Cohorts

#### Autism cohorts

We studied two autism samples and intra-familial controls when available (Figure 1 and Table S1 in the online supplement). The Simons Simplex Collection (SSC) (30), a cohort of 2,569 simplex families: 2,074 quads (one autistic proband, unaffected parents, and one unaffected sibling) and 495 trios (one autistic proband and unaffected parents). The MSSNG database, used as an independent replication cohort, includes 1,381 probands with autism. (31)

#### Unselected cohorts

We included 2,769 individuals from two community-based cohorts that we previously studied (26): IMAGEN (N=1,802) (32) and the Saguenay Youth Study (SYS; N=967) (33) (Figure 1 and Table S1 in the online supplement).

### CNV calling and annotation

We analyzed genotyping data from SSC, IMAGEN, and SYS and whole genome sequencing data from MSSNG. CNV detection, filtering, and annotation are detailed in Methods in the online supplement. We attributed 9 scores to deletions and duplications. These included size, number of genes, number of expression quantitative trait loci regulating genes expressed in the brain (34). Each coding gene with all isoforms fully encompassed in CNVs was annotated using 4 constraint scores which reflect genetic fitness: the pLI score (ExAC v1.0), which is available for 18,224 genes and ranges from 0 – meaning that the gene is tolerant to mutations – to 1 – the gene is intolerant to mutations (27), the residual variation intolerance score (RVIS) (35), the deletion and duplication scores from ExAC (36). Coding genes were also scored using the number of protein-protein interactions (37) and the differential stability score (38). We computed the ancestry in the SSC, IMAGEN and SYS cohorts based on HapMap3 reference population. (39)

### Clinical assessments

NVIQ data were available across all cohorts. (30–33) The assessment methods are detailed in Methods and Table S2 in the online supplement. All other cognitive, behavioural, and motor phenotypes are detailed in Table 1, Methods and Table S1 in the online supplement. Participants underwent age- and development-appropriate standardized cognitive and behavioural tests.

### Statistical analyses

#### Effect-size of gene dosage on general intelligence in probands and the unselected populations

For each individual, we computed the sum of a given score for deletions and duplications separately (Figure 1, Methods in the online supplement). These deletion and duplication scores were used as two independent main effects in the model. We performed a stepwise variable selection procedure based on Bayesian information criteria to identify which score (among the 9 tested) best explain NVIQ for deletions and duplications. This was performed independently for the SSC probands, the unselected populations, and MSSNG as a replication dataset. Age, sex, ancestry and familial relatedness were used as covariates when applicable (Methods in the online supplement).

#### Effect-size of gene dosage on autism risk

We performed the same stepwise variable selection procedure to identify CNV scores that best explain the effect-size of deletions and duplications on autism risk. The dependent variable was the binary diagnosis (autism/control) and independent variables were the selected CNV scores. Conditional logistic regression was used when matching SSC probands with their unaffected siblings. Simple logistic regression was used when comparing SSC probands with the unselected populations. We assessed the effect-size of gene dosage on autism risk beyond its effect on NVIQ by adjusting for NVIQ or performing 1:1 matching of probands with individuals from the unselected populations based on NVIQ (Methods and Figure S1D in the online supplement). Replication analyses were performed using the MSSNG dataset. Sex, ancestry and familial relatedness were used as covariates when applicable (Methods in the online supplement).

To estimate the proportion of autism-risk potentially mediated by NVIQ for deletions and duplication, we performed a counterfactual-based mediation analysis on the pooled dataset.

#### Sensitivity analyses

For sensitivity analyses, we pooled all samples and excluded individuals with CNVs > 10 points of pLI (deletions with an effect > 2 standard deviations of NVIQ) or recurrent CNVs associated with neurodevelopmental disorders or rare *de-novo* CNVs (Tables S3, S4 and S5 in the online supplement).

#### Estimating and validating the level of autism risk

We compared the autism risk estimated by our model to that previously published for recurrent CNVs. Our literature search identified 16 CNVs with available odds ratios (ORs) (9, 12, 13, 40) (Table S6 in the online supplement). The model was trained using a pooled dataset including SSC and MSSNG probands, unaffected siblings, and unselected populations, excluding these 16 CNVs.

To illustrate the output of our model, we computed the autism risk for each CNV called in both autism cohorts including at least one gene with a pLI annotation. We also computed autism risk for any 1MB CNV across the genome, generating a series of 1Mb deletions and duplications (Human Gene Nomenclature) by moving a sliding window in 50Kb steps across the genome. (41) We chose 1Mb CNVs based on thresholds for deleteriousness used in previous studies. (23, 42)

#### Effect-size of gene dosage on measures of core symptoms and specifiers of autism

We investigated the effect of the previously selected CNVs score on cognitive, behavioural, and motor phenotypes to understand why they increase susceptibility to autism. The choice of the statistical model depended on the distribution of the phenotypic measure (Methods and Table S7 in the online supplement). The Social Responsiveness Scale (SRS) was investigated using the entire SSC, MSSNG probands and IMAGEN cohorts (Methods and Table S8 in the online supplement). The Autism Diagnostic Observation Schedule (ADOS) and Autism Diagnostic Interview-Revised (ADI-R) were investigated using probands from SSC and MSSNG (Methods and Table S9 in the online supplements). The Child behaviour Checklist (CBCL) was investigated on probands and unaffected siblings from the SSC (Methods and Table S10 in the online supplements). All other phenotypic measurements were analysed using SSC probands alone. For all analyses, age, sex, ancestry and familial relatedness were used as covariates when applicable. Phenotypic measures were also tested with and without adjustment for NVIQ and/or autism diagnosis when available (Methods in the online supplement). Computation of the significance threshold is detailed in Methods in the online supplement.

## RESULTS

### Effect-size of gene dosage on general intelligence in probands and the unselected populations

As we previously observed in unselected populations (26), the variable selection procedure identified the sum of pLI scores as the variable that best explains the variance of NVIQ in the SSC for deletions (r^2^=0.013) and duplications (r^2^=0.004), compared to the 8 other scores. The sum of pLI scores per individual ranges from 0 to 18.92 and 35.71 for deletions and duplications respectively.

A point of pLI – which correspond to the score of a gene intolerant to mutations – in deletion has the same effect-size on z-scored NVIQ in autism probands of both samples (SSC: β=−0.17, SE=0.03, p=8×10^−10^; MSSNG: β=−0.20, SE=0.07, p=3×10^−3^) and unselected populations (β=−0.19, SE=0.04, p=7×10^−5^). The pLI is also the score that best explains the impact of duplications on NVIQ, showing a three-fold smaller effect of pLI points on z-scored NVIQ in the SSC (β=−0.06, SE=0.02, p=1×10^−3^). No significant effect of duplications is detected in unselected populations or the MSSNG dataset (Table S11 in the online supplement, Figure 2A).

**Figure 2.**
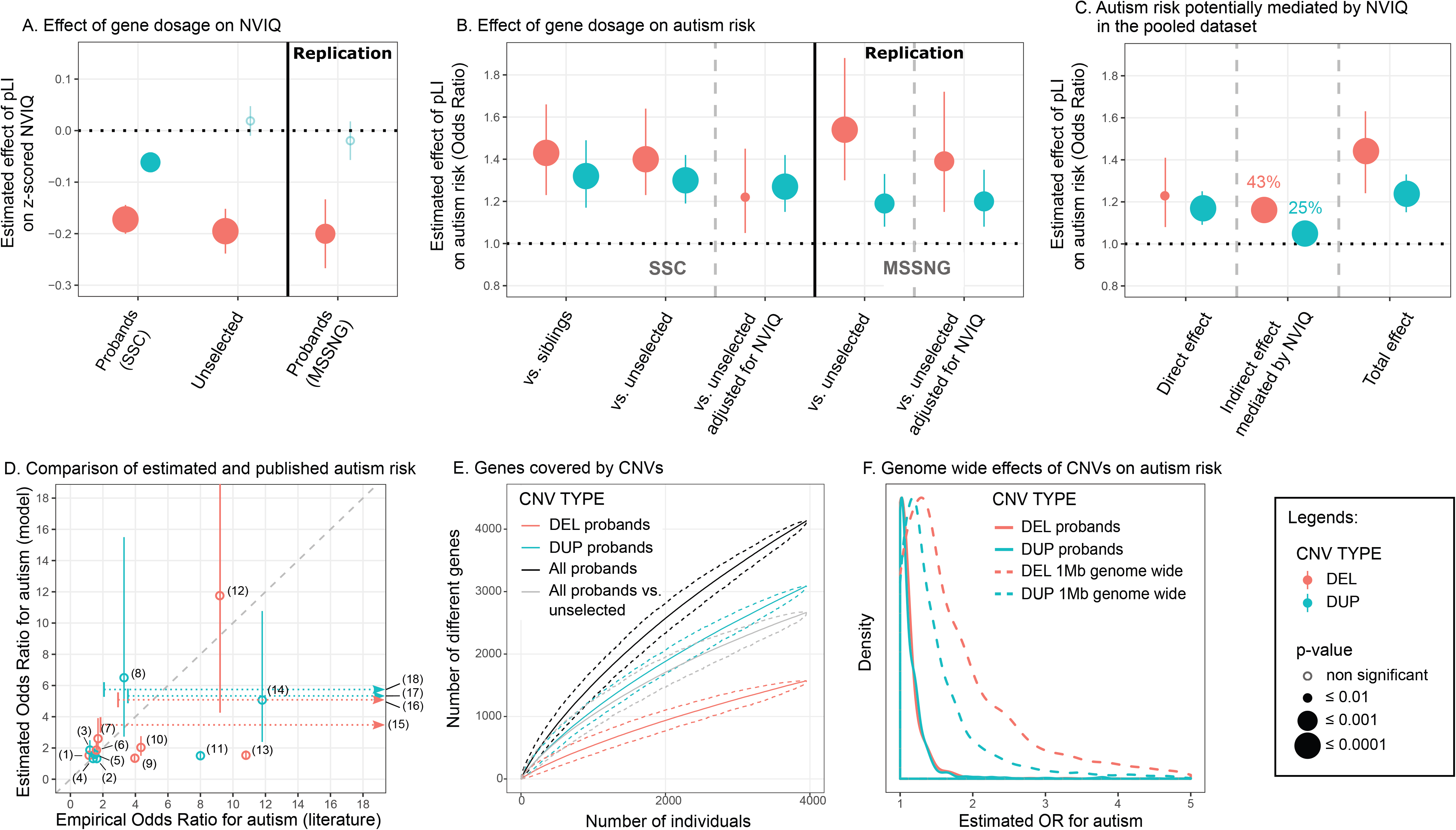
Effect of gene dosage on NVIQ and autism susceptibility. (A) Effect-size of a deleted (DEL, red) or duplicated (DUP, blue) point of pLI on NVIQ in autistic probands and unselected populations. Y axis values are z-scores for NVIQ (e.g. 0.2 z-score=3 points of NVIQ). (B) Autism risk conferred by a deleted (red) or duplicated (blue) point of pLI. Y axis values are odds ratios (OR) computed using a logistic regression to explain an autism diagnosis. Controls include unaffected siblings or unselected populations. Replication was performed using autistic probands from the MSSNG dataset and the unselected populations. (C) Autism risk potentially mediated by NVIQ on the pooled dataset. Y axis values are ORs computed using a counterfactual-based mediation analysis on a logistic regression. Direct effects of CNVs are those not mediated by NVIQ. Indirect are those potentially mediated by NVIQ. Percentage of effect mediated by NVIQ=(indirect/total effect)×100. Total effects are those computed without adjusting for NVIQ. (D) We compared the risk of autism estimated by our model and the risk observed in previous published studies on 16 recurrent CNVs (4, 9, 22) (Table S3 in the online supplement). OR estimates from the model overlap with the 95% confidence interval (CI) of ORs from previous publications for 14 recurrent CNVs. For three CNVs, the horizontal dotted arrows represent the extreme variability for ORs reported in previous publications. Values are detailed in Table S3 in the online supplement. (1) DEL 15q11.2, (2) DUP 15q11.2, (3) DUP 16p11.2 distal, (4) DUP 15q13.3, (5) DUP 16p13.11, (6) DEL 1q21.1, (7) DEL 16p11.2 distal, (8) DUP 22q11.2, (9) DEL 17p12, (10) DEL 16p13.11, (11) DUP 1q21.1, (12) DEL 16p11.2, (13) DEL 15q13.3, (14) DUP 16p11.2, (15) DEL 17q12, (16) DEL 3q29, (17) DUP 7q11.23 and (18) DUP 17p11.2. (E) Genes covered by deletions (red), duplications (blue), or both (black) in autistic probands. The grey line represents the excess of genes encompassed in CNVs in autistic probands relative to the unselected populations. The Y-axis represents the number of distinct genes encompassed in CNVs (each gene is only counted once). The X-axis represents the number of individuals. The mean and 95%CI confidence interval were obtained using 1,000 iterations (bootstrap procedure). (F) Distribution of the estimated effects of deletions (red) and duplications (blue). Full line: Estimated effects computed for all CNVs ≥ 50Kb identified in both autism cohorts. Dotted line: Autism risk computed for any CNV of 1MB across the genome including at least one gene with a pLI annotation.

In the pooled dataset, an autism diagnosis does not influence the effect of deleted or duplicated points of pLI on NVIQ. There is also no interaction with sex. Removing carriers of CNVs with a pLI sum > 10, with a known psychiatric association, or one occurring *de-novo*, results in similar effect-sizes for deletions. For duplications, our limited power only allowed us to observe an effect when removing CNVs enriched in neurodevelopmental disorders (Table S4 in the online supplement).

### Effect-size of gene dosage on autism risk

The variable selection procedure identified again the sum of pLI scores as the variable that best explains the diagnosis of autism for deletions (r^2^=0.004) and duplications (r^2^=0.004). Susceptibility to autism significantly increases for each deleted point of pLI and the effect-size is identical when comparing autistic probands with their paired siblings or unselected populations (OR=1.43, 95%CI=1.23-1.66, p=4×10^−6^; OR=1.40, 95%CI=1.23-1.64, p=2×10^−6^, respectively). A duplicated point of pLI also significantly increases autism susceptibility (comparing with siblings: OR=1.32, 95%CI=1.17-1.49, p=5×10^−6^; and the unselected populations: OR=1.30, 95%CI=1.19-1.42, p=2×10^−8^) (Figure 2B, Table S12 in the online supplement). Of note, there is no difference in pLI burden between intra- and extra-familial controls (unselected populations) (Table S5 in the online supplement).

The risk conferred by deletions measured by pLI decreases substantially but remains borderline significant when the model is adjusted for NVIQ (OR=1.22, 95%CI=1.05-1.45, p=0.01) or when both autism and unselected populations are matched for NVIQ. In contrast, the autism risk conferred by each duplicated point of pLI remains unchanged when adjusting (OR=1.27, 95%CI=1.15-1.42, p=5×10^−6^) or matching for NVIQ (Figure 2B and Table S12 in the online supplement).

The replication analysis with the MSSNG dataset shows the same effect of deleted or duplicated points of pLI on autism susceptibility. We also replicate the differential effect of NVIQ adjustment on autism risk conferred by deletions and duplications (Figure 2B and Table S12 in the online supplement).

In the pooled dataset mediation analysis suggested that 43% and 25% of the autism risk conferred by deletions and duplications are potentially influenced by NVIQ (Figure 2C and Table S13). However, the effect-size of autism risk for deletions and duplications measured by pLI is the same and is significant in both subgroups of individuals above and below median NVIQ (Figure S2 and Table S14). There is no interaction with sex. Autism susceptibility related to gene dosage is unaffected by removing carriers of CNVs with a pLI sum > 10, CNVs with a known association to neurodevelopmental disorder, occurring *de-novo,* or individuals from the unselected populations with a suspected diagnosis of autism (n=10) as well as no diagnostic information from the Development and Well-Being Assessment (DAWBA) (N=124) (Table S5 in the online supplement).

### Estimating and validating the level of autism risk

ORs have previously been computed for a few recurrent CNVs with broad confidence intervals. The autism risk estimated by our model overlaps with that previously published for 16 recurrent CNVs, except for the deletion 15q13.3 and the duplication 16p11.2 which are discordant (9, 12, 13, 40) (Figure 2D, Table S6 in the online supplement). The results are similar whether we include or exclude, the 16 CNVs from the training dataset (Figure S3 in the online supplement). Our model is trained on deletions and duplications covering over 4,500 different genes in the autism and unselected populations (Figure 2E). The sharply ascending slope of genes encompassed in the CNVs shows no asymptotic effects. Model estimates show that any 1Mb coding deletion or duplication across the genome should increase autism susceptibility, with a median OR of 1.6 and 1.3, respectively (Figure 2F and Table S15 in the online supplement).

### Effect-size of gene dosage on measures of core symptoms and specifiers of autism

We assessed the cognitive and behavioural symptoms that underlie autism susceptibility conferred by gene dosage.

#### Autism related symptoms

The pLI significantly increases the SRS, with a 2:1 effect-size ratio for deletions and duplications in the pooled SSC and IMAGEN dataset (deletions: β=3.72 points of raw SRS score per point of pLI,, SE=0.57, p=5×10^−11^; duplications: β=1.87 points of raw SRS score per point of pLI, SE=0.43, p=1×10^−5^). The effect-size of pLI on SRS remains the same after adding data from MSSNG (deletions: β=3.68, SE=0.56, p=4×10^−11^; duplications: β=1.63, SE=0.42, p=1×10^−4^). This effect of gene dosage is entirely explained by NVIQ and the autism diagnosis (Figure 3A, Table S8 in the online supplement).

**Figure 3.**
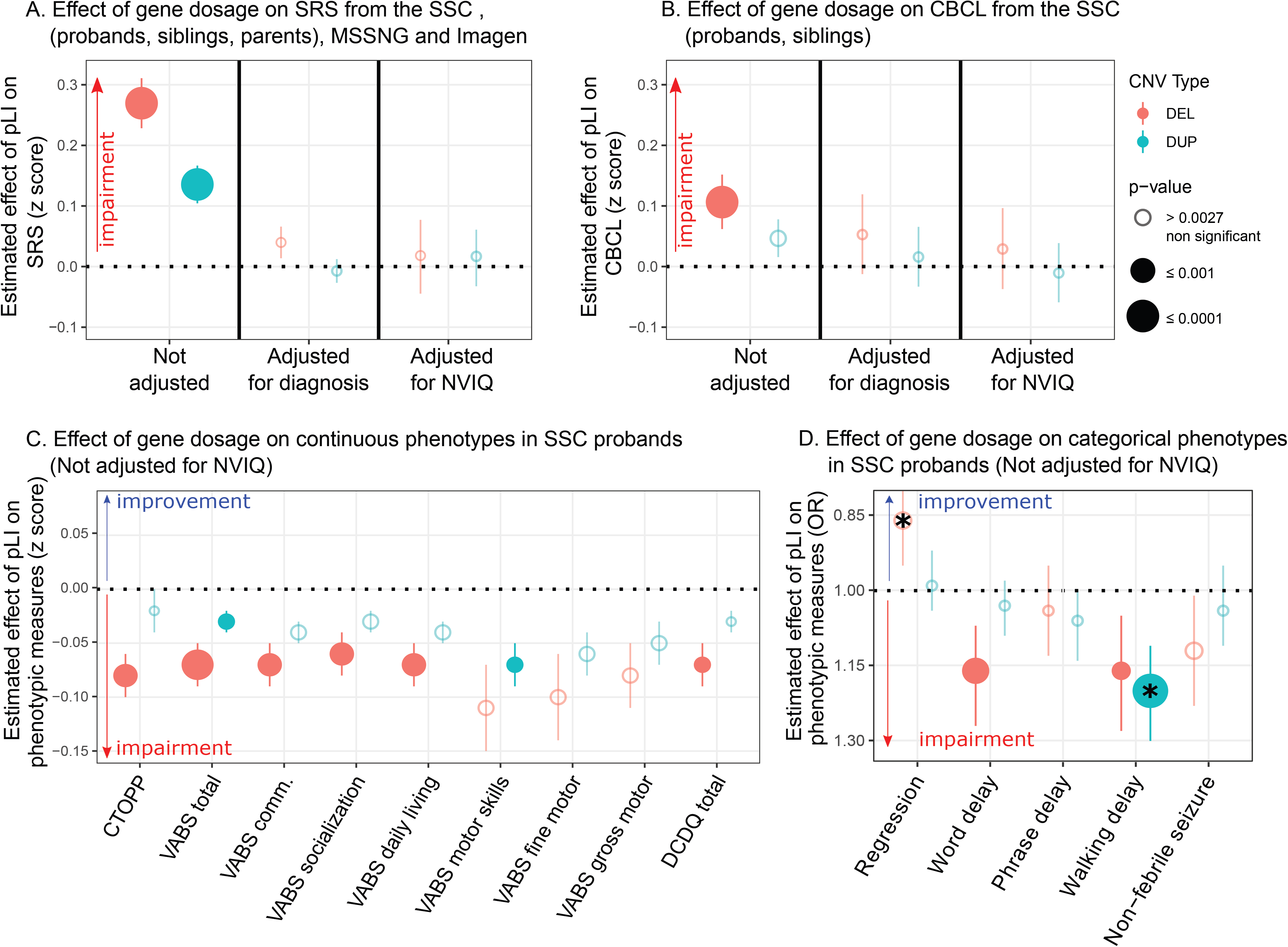
Effect of gene dosage on phenotypic measurements. (A) Effect-size of a deleted (red) or duplicated (blue) point of pLI on the z-scored Social Responsiveness Scale (SRS) in autistic probands from the SSC pooled with their unaffected relatives (siblings and parents) and the unselected populations from the IMAGEN dataset. Effects were measured with and without adjustment for the diagnosis of autism, and NVIQ. The Y-axis represents the estimated effect of pLI on the SRS z-score computed using the mean and standard deviation of unaffected individuals (0.10 z-score = 1.28 SRS raw score point). The analysis adjusting for NVIQ only contains probands from the SSC and individuals from Imagen. (B) Effect-size of a deleted or duplicated point of pLI on the z-score of the Child behaviour Checklist (CBCL) in autistic probands from the SSC pooled with their unaffected siblings. Effects were measured with and without adjustment for the diagnosis of autism, and NVIQ. The estimates were originally computed as OR using a negative binomial regression. The Y-axis represents the estimated effect of pLI on z-scored CBCL computed using the mean and standard deviation of unaffected individuals (0.10 z-score = 1.52 CBCL raw score point). The analysis adjusting for NVIQ is only performed on probands from the SSC. (C, D) Effect-size of a deleted (red) or duplicated (blue) point of pLI on continuous (C) and categorical (D) phenotypes in autistic probands from the SSC unadjusted for NVIQ. Results adjusted for NVIQ are detailed in Figure S4. Y-axis values of panel (C) are measures z-scored using normative data (Table S7) except for DCDQ which was z-scored using the SSC autistic proband group. Y-axis values of panel (D) are odds ratios computed by logistic regression. The significance threshold was computed (51) to account for multiple testing: 0.0027. * Effects significant after adjusting for NVIQ. CTOPP: Comprehensive Test of Phonological Processing; VABS-II: Vineland Adaptive behaviour Rating Scales - Second Edition; DCDQ: Developmental Coordination Disorder Questionnaire.

Deletions and duplications measured by pLI do not affect the ADOS or ADI-R scores in probands of the SSC and MSSNG datasets, pooled or separately (Table S9 in the online supplement). Moreover, deletions measured by pLI protect against regression in autism and this effect is enhanced after adjusting for NVIQ (OR=0.80, 95%CI=0.70-0.89, p=2×10^−4^) (Table 1, Figure 3D, Figure S4B in the online supplement).

#### Language and phonological memory

There is a dissociation between the effect of deletions and duplications on language. Deleted points of pLI are associated with a delay of first-words (OR=1.16, 95%CI=1.07-1.27, p=5×10^−4^) and negatively affects phonological memory, assessed by the non-word repetition of the Comprehensive Test of Phonological Processing (CTOPP) (β=0.08, SE=0.02, p=6×10^−4^). No effects are observed for duplications or after adjusting for NVIQ (Table 1, Figure 3C and 3D, Figure S4A and 3B in the online supplement).

#### Behavioural and emotional symptoms

In the sample pooling probands and unaffected siblings, haploinsufficiency measured by pLI impacts the score of total problems from the CBCL (OR=1.05, 95%CI=1.03-1.08, p=2×10^−6^). The effect of duplications is weaker (OR=1.02, 95%CI=1.01-1.04, p=3×10^−3^) (Table S10, Figure 3B). This translates into an increase of 20.63 [95%CI=19.55-21.73] and 7.85 [95%CI=7.28-8.44] points for a deletion or a duplication encompassing 10 points of pLI, respectively. These effects are not observed within SSC probands or unaffected siblings samples.

#### Adaptive Skills

Adaptive skills measured by the second edition of the Vineland Adaptive behaviour Rating Scales (VABS-II) are negatively affected by the pLI, with a decrease of 2 and 1 point of VABS per deleted or duplicated point of pLI, respectively (p=3×10^−5^ and p=3×10^−3^). Total scores and all subscales are equally affected. NVIQ appears to account for most, if not all, of this effect (Table 1, Figure 3C, Figure S4A in the online supplement).

#### Motor skills and epilepsy

The relationship between the onset of walking measured in months and pLI (deletion: OR=1.03, 95%CI=1.02-1.04, p=2×10^−11^; duplication: OR=1.02, 95%CI=1.01-1.03, p=7×10^−9^) translates into a 5.46 [95%CI=5.27-5.65] or 3.58 [95%CI=3.45-3.72] month delay for a deletion or duplication encompassing 10 points of pLI, respectively (Figure S5 in the online supplement). This remains significant after adjusting for NVIQ for duplications only. The effect-size of gene dosage on motor skills, measured by the VABS-II and the Developmental Coordination Disorder Questionnaire (DCDQ), shows a 2:1 ratio for deletions and duplications with a similar effect for gross and fine motor skills. Gene dosage does not affect the risk of non-febrile seizures (Table 1, Figure 3C and 3D, Figure S4 A and B in the online supplement).

### Potential applications in the clinic

We developed a prediction tool available online (https://cnvprediction.urca.ca/) to estimate the effect-size of deletions and duplications on NVIQ, autism risk and the SRS score. As an illustration, our model estimates a decrease in NVIQ of 26.78 [95%CI=26.19-27.37] and 30.89 [95%CI=30.30-31.48] points, an increase in the SRS raw score of 36.93 [95%CI=35.82-38.04] and 42.59 [95%CI=41.48-43.70] points, and an increase in autism risk of 21.05 [95%CI=6.10-72.26] and 33.58 [95%CI=8.05-139.99] for the 16p11.2 and 22q11.2 deletion respectively. We detail the model output for 21 recurrent CNVs in Table S6. Briefly summarized, this tool should be viewed as a translation of gnomAD (43) information into phenotypic effect-sizes.

## DISCUSSION

We propose a model to estimate the effect-size of gene dosage on autism susceptibility, core autism symptoms, general intelligence, and autism specifiers. Haploinsufficiency measured by pLI increases autism susceptibility across the genome but NVIQ drives a large proportion of this effect. Language, motor, social communication, and behavioural problems are also strongly affected by deletions. While these manifestations may increase the probability for deletion carriers of receiving an autism diagnosis, there are no evidence that core symptoms are affected (Figure 4). In contrast, duplicated points of pLI increase autism risk, genome wide, and the influence of NVIQ is smaller. Increased risk measured by pLI is significant and similar in subgroups of individuals with NVIQ below and above median.

**Figure 4.**
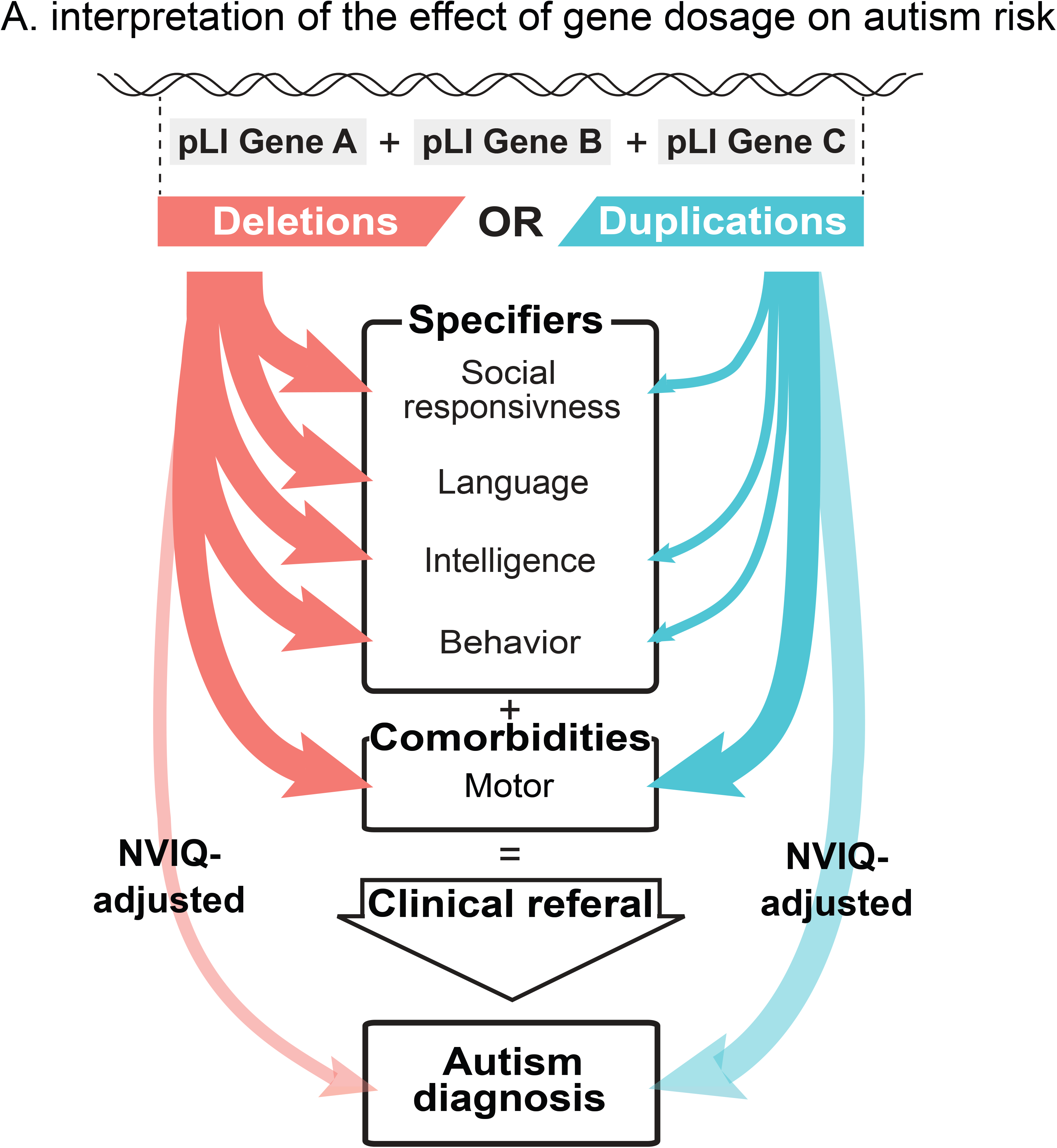
Interpretation of the effect of gene dosage on autism risk. Summary interpretation of the differential effects of deletions and duplications on autism risk. Effect sizes of deletion on most autism specifiers are larger than those of duplications. Duplications and to a lesser extent deletions increase the probability of an autism diagnosis after adjusting for their effect on NVIQ (transparent arrows).

### Differential effects of deletions and duplications on autism core symptoms and specifiers

Model estimates show that any 1Mb coding deletion or duplication across the genome should increase autism susceptibility, with a median OR of 1.6 and 1.3, respectively (Figure 4B). GWAS conducted on common variants also showed that the bulk of the heritability for complex conditions (*i.e.* schizophrenia) is spread across the genome and largely driven by genes with no clear relevance to disease. (29, 44) Gene dosage affects NVIQ, social communication, and adaptive behaviour, with a deletion:duplication effect-size ratio of 2-3:1. Although both CNVs equally affect motor skills, phonological memory may be predominantly affected by haploinsufficiency. Similar differential profiles have been reported for 16p11.2 CNVs with phonological memory deficits in deletion but not duplication carriers. (45) The phenotypic profile of gene dosage may apply across the genome, independently of effect-size. The phenotypic profile of haploinsufficiency delineated by our model has been similarly reported in patients with *de-novo* loss of function variants (24, 25). In addition, excluding large effect-size *de-novo* variants from our analyses does not modify effect of gene dosage, measured by pLI, on NVIQ and autism risk. Therefore, molecular functional networks enriched in genes with an excess of *de-novo* mutations (chromatin remodelling, synaptic function) (15, 46, 47) may be related to large effect-sizes rather than specific effects on autism risk. Interestingly, the lack of interaction between the effect of pLI and sex suggests that a point of pLI similarly affects NVIQ and increases autism risk in both sexes.

### Clinical applications

Our models are implemented in a prediction tool (https://cnvprediction.urca.ca/), which is designed to predict the effect-size of CNVs, not the symptoms of the individual who carries the CNV. If the symptom severity of the individual is concordant with the effect-size of the CNV, we may conclude that the CNV contributes substantially to the clinical phenotype. If symptoms are discordant, the clinician may conclude that additional factors should be investigated. The estimates of autism risk provided by models in this study overlap with risk computed in previous studies. As an example, our model estimates for 16p11.2 and 22q11.2 deletions are similar to the previously published effect for NVIQ, (loss of 25 (48) and 29 (49) points), autism risk (OR of 11.8 (12) and 32.37 (9)) and SRS (gain of 44 (48) and 49 (49) points). Overall, the output of these models can help interpret CNVs in the clinic, but these mean estimates may not apply to many CNVs.

### Limitations

Discordance between autism risk estimated by the model and literature observations allows for the identification of CNVs, which may encompass genes with specific properties. For example, autism susceptibility and deficits associated with the 15q13.3 (*CHRNA7*) deletion appear to be underestimated by our model. This CNV may include genes for which the assigned pLI score does not capture the effects on psychiatric traits (*e.g.* gene dosage of *CHRNA7*, which has a pLI=0 may affect psychopathology without altering genetic fitness). The pLI was not developed to measure intolerance to duplications and results should, therefore, be interpreted with caution. Our findings suggest, however that pLI may be a general measure of dosage sensitivity, in line with recent data from gnomAD-SV. (50) Since gene dosage is not comparable between sex-linked and autosomal CNVs, we could not pool both types of CNVs. Sex-linked CNVs were excluded from this study because they were too rare in our samples to be studied separately. The effect of gene dosage on SRS was very robust but was mainly explained by the autism diagnosis. This suggests that the SRS may not measure a continuous dimension since this score is unable to provide additional granularity within the autism group or the controls despite large sample size. Some phenotypic measures such as phonological memory and motor skills were only available for autism probands and results may not be generalizable to non-autism samples. Larger samples, with additional intrafamilial controls, novel functional annotations, and more refined models are required to improve our estimates of CNV effect-sizes on cognitive dimensions.

## Conclusion

Our study highlights the extreme polygenicity of autism susceptibility conferred by gene dosage. It also delineates cognitive mechanisms which may explain in part the overrepresentation of CNVs in autism. Among mutations over-represented in autism, those truly related to core symptoms may be less common than previously thought. Future large-scale studies simultaneously investigating the effect of genomic variants on categorical diagnoses and continuous dimensions are warranted. This study represents a new framework to study rare variants and can help in the interpretation of the effect-size of undocumented CNVs identified in the neurodevelopmental clinic.

## Supporting information

Supplemental content

## Funding/Support

This research was enabled by support provided by Calcul Quebec (http://www.calculquebec.ca) and Compute Canada (http://www.computecanada.ca). Dr. Jacquemont is a recipient of a Bursary Professor fellowship of the Swiss National Science Foundation, a Canada Research Chair in neurodevelopmental disorders, and a chair from the Jeanne et Jean Louis Levesque Foundation. Dr. Schramm is supported by an Institute for Data Valorization (IVADO) fellowship. Ms. Tamer is supported by a Canadian Institute of Health Research (CIHR) Scholarship Program. Dr. Huguet is supported by the Sainte-Justine Foundation, the Merit Scholarship Program for foreign students, and the Network of Applied Genetic Medicine fellowships. Dr. Loth is supported by European Autism Interventions, which receives support from the Innovative Medicines Initiative Joint Undertaking under grant agreement 115300, the resources of which are composed of financial contributions from grant FP7/2007-2013 from the European Union’s Seventh Framework Programme, the European Federation of Pharmaceutical Industries and Associations companies’ in-kind contributions, and Autism Speaks. Dr. Bourgeron is a recipient of a chair of the Bettencourt-Schueler foundation. Laurent Mottron is a recipient of the Marcel & Rolande Gosselin research chair. This work is supported by a grant from the Brain Canada Multi-Investigator initiative and CIHR grant 159734 (Jacquemont, Greenwood, Paus). The Canadian Institutes of Health Research and the Heart and Stroke Foundation of Canada fund the Saguenay Youth Study. SYS was funded by the Canadian Institutes of Health Research (TP, ZP) and the Heart and Stroke Foundation of Canada (ZP). Funding for the project was provided by the Wellcome Trust. This work was also supported by an NIH award U01 MH119690 granted to Drs. Almasy, Jacquemont and Glahn and U01 MH119739. The authors gratefully acknowledge the resources provided by the Autism Speaks MSSNG project and the Autism Genetic Resource Exchange Consortium, as well as the participating families.

We are grateful to all the families who participated in the Simons Variation in Individuals Project (VIP) and the Simons VIP Consortium (data from Simons VIP are available through SFARI Base). We thank the coordinators and staff at the Simons VIP and SCC sites. We are grateful to all of the families at the participating SSC sites and the principal investigators (A. Beaudet, M.D., R. Bernier, Ph.D., J. Constantino, M.D., E. Cook, M.D., E. Fombonne, M.D., D. Geschwind, M.D., Ph.D., R. Goin-Kochel, Ph.D., E. Hanson, Ph.D., D. Grice, M.D., A. Klin, Ph.D., D. Ledbetter, Ph.D., C. Lord, Ph.D., C. Martin, Ph.D., D. Martin, M.D., Ph.D., R. Maxim, M.D., J. Miles, M.D., Ph.D., O. Ousley, Ph.D., K. Pelphrey, Ph.D., B. Peterson, M.D., J. Piggot, M.D., C. Saulnier, Ph.D., M. State, M.D., Ph.D., W. Stone, Ph.D., J. Sutcliffe, Ph.D., C. Walsh, M.D., Ph.D., Z. Warren, Ph.D., and E. Wijsman, Ph.D.). We appreciate obtaining access to phenotypic data on SFARI base.

## Additional Contributions

Julien Buratti (Institute Pasteur), and Vincent Frouin, Ph.D. (Neurospin), acquired data for IMAGEN. Manon Bernard, BSc (database architect, The Hospital for Sick Children), and Helene Simard, MA, and her team of research assistants (Cégep de Jonquière) acquired data for the Saguenay Youth Study. Antoine Main, M.Sc. (UHC Sainte-Justine Research Center, HEC Montreal), Lionel Lemogo, M.Sc. (UHC Sainte-Justine Research Center), and Claudine Passo, Pg.D. (UHC Sainte-Justine Research Center), provided bioinformatical support. Maude Auger, Pg.D. (UHC Sainte-Justine Research Center), provided website development. Dr. Paus is the Tanenbaum Chair in Population Neuroscience at the Rotman Research Institute, University of Toronto, and the Dr. John and Consuela Phelan Scholar at Child Mind Institute, New York.

