## Supplemental content for "Effects-sizes of deletions and duplications on autism risk across the genome"

**SUPPLEMENTARY ONLINE CONTENT**

**TABLE OF CONTENTS**

**Methods. Supplemental Methods**

**Figure S1: Distribution of NVIQ.**

**Figure S2: Distribution of NVIQ for individual below or above the median (98), using probands from SSC and MSSNG, and the unselected populations.**

**Figure S3: Effect of gene dosage measured by pLI on autism risk.**

**Figure S4: Effect of gene dosage measured by pLI on phenotypic measures adjusted for NVIQ in SSC probands.**

**Figure S5: Effect of gene dosage measured by pLI on age of onset for walking.**

**Figure S6: Effect of gene dosage measured by pLI on SRS score.**

**Table S1: Description of cohorts: Demographic, genotypical characteristics and phenotypical measures.**

**Table S2: NVIQ available in autistic probands from SSC and MSSNG, and individuals from unselected population.**

**Table S3: Breakpoints used to detect recurrent CNVs associated to neurodevelopmental disorders.**

**Table S4: Sensitivity analysis of the effect of gene dosage measured by pLI on NVIQ in autistic probands from SSC and MSSNG, and in the unselected population.**

**Table S5: Sensitivity analysis of the effect of gene dosage measured by pLI on autism risk using autistic probands from SSC and MSSNG, unaffected siblings from SSC, and the unselected population.**

**Table S6: Breakpoints used to detect the 16 recurrent CNVs from the comparison of published and estimated odds ratios in autistic population.**

**Table S7: Description of models used for the investigation of the effect of gene dosage on phenotypical measures of autistic probands from SSC.**

**Table S8: Effect of gene dosage measured by pLI on SRS using autistic probands from SSC, unaffected siblings, unaffected parents and the unselected population from IMAGEN.**

**Table S9: Effect of gene dosage measured by pLI on severity scores of autism (main domains of ADI-R and ADOS-calibrated severity scores) using autistic probands from SSC and MSSNG.**

**Table S10: Effect of gene dosage measured by pLI on CBCL using autistic probands from SSC and unaffected siblings.**

**Table S11: Effect of gene dosage measured by pLI on general intelligence in autistic probands from SSC and MSSNG, and in the unselected population.**

**Table S12: Effect of gene dosage measured by pLI on autism risk.**

**Table S13: Autism risk potentially mediated by NVIQ in the pooled dataset (SSC, MSSNG, unselected populations).**

**Table S14: Autism risk measured by pLI in subgroups of individual below or above the median (98) in the pooled dataset (SSC, MSSNG, unselected populations).**

**Table S15: Estimated Genome wide effects of gene dosage on autism risk.**

**References.**

**This supplementary material has been provided by the authors to give readers additional information about their work.**

**SUPPLEMENTARY METHODS**

**Definition of autism**
Autism spectrum disorder (ASD) refers to autism as defined in DSM-V (1). As the participants involved in the current study were diagnosed following DSM-IV criteria which uses a subtyping strategy (autistic disorder, Asperger syndrome, and PDDNOS) (2), we will use the generic term ‘’autism” to avoid confusion.

**CNV detection, annotation and filtering**

*Genotyping and whole genome sequencing*

- *Genotyping data*

CNV detections and standard filtering strategies were previously published (3). CNV calling was performed using the same pipeline for individuals from the Simons Simplex Collection (SSC) (4), IMAGEN (5), and Saguenay Youth Study (SYS) (6) to obtain a harmonized dataset.

In the IMAGEN cohort (5), 2,090 individuals were genotyped using a combination of the Illumina 610Kq (N probes=620,901; N arrays=708) and 660Wq (N probes=657,366; N arrays=1,385). The genotyping was performed at the Centre National de Genotypage (CNG; Paris, France).

In SYS cohort (6), 1,994 Illumina SNP arrays were analyzed using Illumina 610Kq (N probes=620,901; N arrays=599) and the HumanOmniExpress BeadChip - V12 (HOE-V12) (N probes=730,525; N arrays=1,395). The genotyping was performed at CNG for 610Kq and at the Genome Analysis Centre of Helmholtz Zentrum München (Munich, Germany) for HOE-V12.

In the SSC (4), 10,032 individuals were genotyped at Yale University using Illumina SNP genotyping arrays 1Mv1, 1Mv3 Duo, or Omni2.5M.

- *Whole genome sequencing data*

In the MSSNG database (7), 7,233 individuals were sequenced at multiple sites using Illumina sequencing HiSeq, HiSeq 2,500, or HiSeqX.

Next generation sequencing data were analysed using Broadinstitute Genome Analysis Toolkit (GATK) best practice (8).

*Call of CNVs*

CNVs from SSC, IMAGEN and SYS were called using PennCNV (9) and QuantiSNP (10) with the following parameters:

- Number of consecutive probes for CNV detection ≥ 3
- CNV size ≥ 1Kb
- Confidence scores ≥ 15.

Then, we merged detected CNVs from both algorithms with CNVision (11).

For MSSNG, read alignment data were used to compute CNV calling following the workflow of Trost et al. (12).

*Filtering of micro-array*

To ensure good quality of CNVs, we kept only micro-arrays without too much noise.

- For IMAGEN and SYS cohorts:
- Wave Factor (WF) < |0.05|
- Standard deviation of the Log-R-Ratio (LRR-SD) < 0.35
- Standard deviation of the B allele frequency (BAF-SD) < 0.08
- Call Rate > 0.99
- For SSC cohort: all micro-array detecting ≥ 200 CNVs were considered as noisy and were removed from the analysis.

*CNV coordinates*

The CNVs coordinates were updated from hg18 to hg19 using Illumina information and the liftover tool from the genome browser (https://genome.ucsc.edu/cgi-bin/hgLiftOver and <http://grch37.ensembl.org/Homo_sapiens/Tools/AssemblyConverter>).

*Concatenation of CNVs*

In a subsequent step, using an in-house algorithm (Pasteur) followed by visual inspection (SnipPeep, http://snippeep.sourceforge.net), we stitched CNVs that appeared to be incorrectly split by the calling algorithms, and we removed any CNVs (size of ≥ 500Kb and ≥ 100 SNPs) that spanned known large assembly gaps (greater than 150Kb).

*CNV filtering*

CNVs with the following criteria were selected for analysis:

- Size ≥ 50 Kb
- Autosomal (Since gene dosage is not comparable between sex-linked and autosomal CNVs, we could not pool both types of CNVs. Sex-linked CNVs were therefore excluded from the analysis.)
- Unambiguous type: deletions or duplications.
- Confidence score ≥ 30 with at least one of both detection algorithms
- Cross array criteria: CNVs overlapping ≥ 10 probes in each of the array technologies used in the study.
- And additional filters were applied for CNVs which are not 40% overlapping with recurrent CNVs from TableS3: Overlap with Segmental duplicates or centromeric regions < 50%.

*CNV annotation and scoring*

The annotation of CNVs was performed with a R package developed by our team. This package was developed using RefSeq (<https://genome.ucsc.edu/>) and ANNOVAR (13) coding and non-coding genes. For these analyses, we excluded the pseudo-genes (UCSC site: https://genome.ucsc.edu/).

Deletions and duplication were annotated for size, number of genes, number of expression quantitative trait loci regulating genes expressed in the brain (14), and each coding gene with all isoforms fully encompassed in CNVs was annotated using 6 constraint scores. For each individual, we computed the sum of these annotations for deletions and for duplications.

Coding genes were annotated using the following constraint scores and transformations:

- the probability of being loss-of-function intolerant (pLI) (15) between 0 and 1 where 1 means that the gene is completely intolerant;
- the residual variation intolerance score (16) between 0 and 100 was transformed with 100-RVIS such as 100 represents the more intolerant;
- the ExAC CNV score for deletions (17) between -2.62 and 3.81 was transformed with deletion score + min(deletion score) such that it becomes between 0 and 6.43 where 6.43 represents the more intolerant.
- the ExAC CNV score for duplication (17) between -2.53 and 2.86 was transformed with duplication score + min(duplication score) such that it becomes between 0 and 5.39 where 5.39 represents the more intolerant.
- the number of protein-protein interactions defined as the number of proteins interacting with each protein coding gene according to STRINGs Protein v10 for Human database (18) (9606.protein.links.v10.txt.gz; http://string-db.org/) where protein networks were defined based on high confidence (> 0.7) interactions;
- the differential stability (DS) score (19), a correlation-based metric which assess reproducibility of regional patterns of gene expression in the brain between -0.057 and 0.97 was transformed with DS + min(DS) such that it becomes between 0 and 1.027 where a higher score means high specific expression in brain.

For the six scores detailed above, the default value associated to gene without available score was 0.

*De novo CNVs identification*

*De novo* CNVs in the SSC were identified in probands, unaffected siblings, and unselected population from SYS using two previously published datasets (11, 20), combined with our own algorithm developed in R. A CNV was considered as *de novo* only if it was defined as such by all three approaches.

*Frequency*

We used two sources to calculate the frequency. First frequency (DGV frequency) was annotated using the database of genomic variants (DGV) on the basis of a minimal overlap of 70% between the CNV of interest and its more closely resembling CNV displayed in DGV (DGV hg19, <http://www.dgv.tcag.ca>). If CNV was seen in several cohorts, we selected the maximal frequency conditionally to the fact that the sample size of the DGV cohort is ≥ 100; otherwise, frequency was considered as null.

Second frequency was computed on the CNVs of 1,804 adolescents in IMAGEN and 893 parents in SYS from the general population cohort (GP-Cohort frequency). It was computed on the basis of a minimal overlap of 70% between CNVs. Since CNVs with a frequency < 0.1% were visualized to ensure the CNVs veracity, the GP-Cohort frequency was recomputed after excluding false positive CNVs. This process was done iteratively until no more CNV is excluded. Then the CNVs of the entire database (SSC and general population cohorts) were annotated with GP-Cohort frequency on the basis of a minimal overlap of 70% between the CNV of interest and its more closely resembling CNV displayed in GP-Cohort. The frequency of a CNV that is not seen in GP-Cohort was considered as null.

*Definition of a rare CNV*

Throughout the paper, a rare CNV is either a known recurrent CNV (Table S3) with a DGV frequency < 0.1% or a non-recurrent CNVs with the following characteristics: (i) DGV frequency < 0.1%; (ii) < 50% of the CNV is contained in regions present at > 1% in DGV (11, 21, 22); (iii) unselected population frequency 1/1000. All CNVs annotated as rare were manually curated by visual inspection (SnipPeep, <http://snippeep.sourceforge.net>) and false positives were excluded.

*Genetic analysis of pairwise ancestry and population stratification*

Classical multidimensional scaling (MDS) was used to identify ancestry based on the identity by state (IBS) matrices of genetic distances (D) in the IMAGEN, the SYS and the SSC cohorts, based on the reference population HapMap3 (23) with 993 individuals (Hapmap Consortium 2003 (www.hapmap.ncbi.nlm.nih.gov). PLINK (24) ([pngu.mgh.harvard.edu/purcell/plink/](http://pngu.mgh.harvard.edu/purcell/plink/)) was used to do these calculations. SNPs were filtered to keep only autosomal SNPs with minor allele frequency (MAF) > 5% and with good quality, significance threshold for a test of Hardy-Weinberg equilibrium < 1.10^-6^ and missing genotype rates < 10%. Related individuals were identified based on *D* defined by the following formula:

$$D= \frac{IBS2+0.5 IBS1}{N SNP pairs}$$

with *IBS1* and *IBS2* being the number of loci at which a pair of individuals share either 1 or 2 alleles identical by state, respectively, and *N SNP pairs* is the number of loci tested. Pairs of individuals were defined as related when *D* ≥ 0.8.

Ancestry was estimated using Admixture (25) (http://www.genetics.ucla.edu/software/admixture) with reference populations from HapMap3 (23) allowing for 4 ancestry components (Africa, Asia, European and India). Results show a strong European ancestry component in the three datasets. We then performed a principal components analysis based on the variance-standardized relationship matrix and displayed the 3 first ancestry dimensions and associated eigenvalues.

**Clinical assessments**

*Non-verbal Intelligence Quotient (IQ)*

Intellectual ability was measured using standardized tests according to the cognitive level of the participant (Table S2).

For SSC autistic probands, non-verbal intelligence quotient (NVIQ) scores were obtained from the Differential Ability Scales, 2nd Edition (DAS-II) (26) for early years (N=1,031) and school age children (N=1,213), the Wechsler Intelligence Scale for Children, 4th Edition (WISC-IV) (27) (N=45), the Wechsler Abbreviated Scale of Intelligence – First Edition (WASI-I) (28) (N=63) or the Mullen Scales of Early Learning (MSEL) (29) (N=213) (see density distribution of NVIQ by test used in Figure1B). Norm-referenced standard scores (deviation NVIQ) were available for most of the participants (85.10%). However, for individuals from SSC who were not able to obtain a deviation NVIQ due to their age and/or developmental level, ratio IQ were derived by dividing mental age by chronological age and multiplying by 100. See Bishop et al., 2011 for more details concerning convergence between ratio and deviation NVIQ (30).

For MSSNG autistic probands, NVIQ scores were obtained from the Leiter international performance scale – Original and revised (31, 32) (N=372), the raven progressive matrices (33) (N=214), the Stanford-Binet intelligence scale (N=281), the Wechsler Intelligence Scale for Children – Fourth Edition (WISC-IV) (27) (N=46), the Wechsler Abbreviated Scale of Intelligence – First and Second Editions (WASI-I, WASI-II) (28, 34) (N=338) or the Wechsler Preschool and Primary Scale of Intelligence – Fourth Edition (WPPSI-IV) (35) (N=128) (see density distribution of NVIQ by test used in Figure1C). Deviation NVIQ were available for all participants.

Individuals in IMAGEN undertook the fourth edition of the Wechsler intelligence scale for children (WISC-IV) whereas SYS cohort used the third edition (WISC-III) to assess children. Deviation NVIQ were available for all participants.

As a preeminent test of fluid intelligence, the Raven’s progressive matrices, has been reported as a better assessment of general cognitive abilities in autistic individuals than common NVIQ tests (36), we also investigated the matrices subtests measured in SSC probands using the Differential Ability Scales – second edition (DAS-II) (26).

*Social Responsiveness Scale*

For all the individuals from the SSC and for the unselected population from IMAGEN, severity of social deficits was ascertained with scores from the social responsiveness scale (SRS) (37, 38).

*Severity of autism main domains*

For probands from the SSC and MSSNG, the severity of autism main domains were assessed by domain-calibrated scores from the Autism Diagnostic Observation Schedule (ADOS) (39, 40) and the Autism Diagnostic Interview-Revised (ADI-R) (41).

*Child behaviour Checklist*

For SSC probands and their unaffected siblings, behavioural problems were assessed on the Total Problems Score, as well as the Internalizing and Externalizing domains of the Preschool and School-aged Child behaviour Checklist (CBCL) (42, 43).

*Phenotypes only available for probands from the SSC*

- *Language and phonology*

Phonological short-term memory was measured using scaled scores from the non-word repetition subtest of the Comprehensive Test of Phonological Processing (CTOPP) (44).

Language level was measured using age of first words, age of first phrases and the overall level of expressive language from the ADI-R (45) (question 09, question 10 and question 30 respectively).

Words and phrases delay variables in probands have been derived using both items of age of first words and age of first phrases from the ADI-R score as following:

We considered that autistic probands have word or phrase delay if parents reported an age of first words greater than 24 months on question 9 and the first phrases after 33 months on question 10.
Also, if autistic probands scored 993, 994, or 997 on question 9 or 10, they were considered as delayed in the corresponding variable. At the contrary, if they scored of 996 on question 9 or 10, they were considered as non-delayed in the corresponding variable.

- *Adaptative skills*

Standard scores of total adaptive skills, communication, interaction and daily living skills domains were measured using the Vineland Adaptive behaviour Rating Scales - Second Edition (VABS-II). (46)

- *Motor skills*

Motor skills were measured using the corresponding main domain of the VAB-II, as well as gross and fine motor skills subdomains.

We also used the age of onset for walking reported in the ADI-R (question 5) as continuous and categorical measures of gross motor skills. (45)

Walking delay variables in probands have been derived using item of age of onset for walking from the ADI-R score (question 5).

We considered that autistic probands have walking delay if parents reported an age of first walk greater than 18 months on question 5, as well as if they scored 997 at the same item. At the contrary, if they scored of 996 on question 5, they were considered as non-delayed.

Motor coordination was assessed by the Developmental Coordination Disorder Questionnaire (DCDQ).

- *Associated neurological condition*

Finally, the presence of non-febrile seizures was assessed from the ADI-R (question 85) and the Medical History Form.

**Statistical analyses**

*Stepwise variable selection procedure based on Bayesian information criteria*

The stepwise variable selection procedure was used to choose the best model with the smallest Bayesian Information Criterion (BIC). We allowed pairwise interactions conditionally upon main effects to also be included in the model. Since variable selection and model fitting steps were performed on the same dataset, this could induce a selection bias in the estimates. We used a bootstrap (1,000 iterations) estimation of the bias in our final models to correct for this bias. (47)

Genetic variable selection procedure was performed using the boot.stepAIC() function from the R package ‘bootStepAIC’. (48)

*Effect of gene dosage on general intelligence*

The model selected in SSC cohort was applied in the unselected population sample. Both were adjusted for NVIQ test used, sex and ancestry. Moreover, the unselected population sample includes related individuals, thus we included a random effect in the model to take into account the familial relationship.

The model adjusted for familial relationship with a random effect could be written as:
NVIQ ~ 𝜶𝑋+ 𝛾𝑍 + β₁ CNV_DEL_ + β₂ CNV_DUP_

where *X* represents the adjustments covariates (NVIQ test used, sex and ancestry) and *Z* is the familial relatedness; CNV_DUP/DEL_: CNV scoring selected as best genetic explanatory variable in SSC cohort; (α, β₁, β_2_) and 𝛾 are respectively the vectors of coefficients for fixed and random effects. The mixed effect model was computed using lme() function from the ‘nlme’ R package. (49)

- *Replication in MSSNG:*

The model selected in SSC cohort was applied in MSSNG cohort, except that the ancestry is not available for MSSNG dataset and therefore we could not adjust for it. The MSSNG dataset also includes related individuals, thus we included a random effect in the model to take into account the familial relationships.

- *Model with pooled dataset to test the interaction between CNV scoring and diagnosis:*

NVIQ ~ 𝜶𝑋+ 𝛾𝑍 + β₁ CNV_DEL_ + β₂ CNV_DUP +_ β_3_ CNV_DEL_ * diagnosis + β_4_ CNV_DUP_ * diagnosis

where *X* represents the adjustments covariates (NVIQ test used, sex, and diagnosis) and *Z* is the familial relatedness; CNV_DUP/DEL_: CNV scoring selected as best genetic explanatory variable is SSC cohort; (α, β₁, β_2_, β_3_, β_4_) and 𝛾 are respectively the vectors of coefficients for fixed and random effects. The mixed effect model was computed using lme() function from the ‘nlme’ R package. (49)

*Effect of gene dosage on autism risk*

All enrichment analyses were performed by excluding related individuals of MSSNG and general population cohorts, i.e. only one subject by family was included in the analyses.

Conditional logistic regression was performed for the comparison of probands paired to their unaffected siblings using the clogit() function from the R package ‘survival’. (50)

All models for this section included sex and ancestry as covariates when available.

- *Matching of autistic probands and general population regarding the NVIQ*

We used a matching procedure to extract subsets from SSC cohort and the general population to obtain pairs of patient-control (ratio 1:1) with similar NVIQ. Matching was made by searching for the nearest neighbor according to NVIQ only. We allowed a gap of 5 points of NVIQ between patient and its matched control. Matching was performed using the Match() function from the R package ‘Matching’. (51)

The matching procedure depends on initial parameters that are randomly chosen. To avoid the potential error due to randomness, we performed the matching 500 times such that we obtained 500 matched cohorts including 1,411 to 1,469 pairs of patient-control. The Figure1A and 1D represent the distributions of NVIQ in initial cohorts and in an example of matched cohorts.

- *Contribution of NVIQ in autism risk effect*

To estimate the proportion of autism-risk potentially mediated by NVIQ for deletions and duplication, we performed a counterfactual-based mediation analysis on the pooled dataset. We used the neImpute(), neModel(), and neEffdecomp() functions from the R package ‘medflex’ (52) on two logistic regression including sum of pLI in deletions or duplications as the main explanatory variable. We also applied a logistic regression including sum of pLI in deletions and duplications as the two mains explanatory variables to estimate the autism risk conferred by deletions and duplications in both subgroups of individuals above and below median NVIQ.

*Effect of gene dosage on phenotypic measures*

- *SRS as a continuous variable in the SSC probands, unaffected siblings and parents, and in the unselected population from IMAGEN*

We used a linear mixed effect model to quantify the effect of gene dosage measured by pLI scores on SRS total raw score after pooling probands and their unaffected relatives (siblings and parents). A kinship matrix was generated to model the genetic covariance between related individuals using the kinship() function from the R package ‘kinship2’ (53) and this covariance was used as a random effect in the model performed with the function lmekin() from the R package ‘coxme’ (54).

We further explored a potential effect of gene dosage on the SRS within the autism group, unaffected siblings, parents and unselected population from IMAGEN using a linear regression and adjusting for the abnormal distributions with a square root transformation of the SRS scores when necessary (Table S8). All models used were adjusted for age, sex, ancestry, and in a second time for NVIQ and/or for the diagnosis of autism.

- *SRS as a categorical variable in the SSC probands, unaffected siblings and unselected population from IMAGEN*

We also investigated the SRS scores based on the previously published *T*-score categorization (55) as follow:

- ***T*-scores of 76 or higher**: Clinically significant deficits in social functioning that interfere with interactions with others;
- **66 *<T*-scores< 75**: Moderate, signaling some clinically significant social deficits;
- **60 *<T*-scores< 65** : Mild to moderate deficiencies in social behaviour;
- ***T*-scores< 59**: Indicate an individual probably does not have social difficulties indicative of a possible autism diagnosis.

A logistic regression was applied in this pooled dataset (autistic probands; unaffected siblings and unselected population) to investigate the effect of gene dosage on binary categorical SRS: clinical (obtained after merging the moderate, mild and clinically significant categories) and normal (Table S8, Figure S5C and S5D). This logistic regression model took into account the family relatedness as random factor using the glmer() function from the R package ‘lme4’.(56)

A cumulative ordinal regression model was also performed on SRS coding for 4 different levels of social deficits (normal, moderate, mild and clinically significant) (Table S8, Figure S5E and S5F). This model was applied using the function vglm() from the R package ‘VGAM’. (57) All models used were adjusted for age, ancestry, and in a second time for the diagnosis of autism.

- *Autism core symptoms in probands from the SSC and MSSNG*

A cumulative ordinal regression model was used to assess the effect of gene dosage on ADOS and ADI-R domains in autistic probands from the SSC and MSSNG separately, and in the pooled dataset (probands from the SSC and MSSNG).

Ordinal regression (cumulative, parallel slopes):

*ln(P(Y>k)/P(Y<=k)) ~ 𝜶_k_𝑋+β₁ pLI_DEL_ + β₂ pLI_DUP_*

Models assessing the ADI-R were adjusted for age, sex and in a second time for NVIQ. When exploring the ADOS, models were corrected for sex and in a second time for NVIQ (Table S9). Additional correction for the population was applied when the model was run on the pooled dataset.

- *CBCL in probands and unaffected siblings from the SSC*

Early analysis using T scores generated by the CBCL demonstrated that despite T scores being normed for sex and age, sex and age emerged as significant factors affecting scores on the school-aged tests. For all analysis afterwards, raw scores were used and corrected for age and sex in the analysis. Several possible models, including an ordinal model testing binning based on “pre-clinical” and “clinical” thresholds based on T-scores, were discarded with the switch to raw scores. Pre-school and school-aged tests were analyzed separately.

Since raw scores have an oversampling of zeros and are overdispersed, it was necessary to use a negative binomial distribution function to adequately assess the relationship between CBCL scores and gene dosage measured by pLI. Probands and unaffected siblings were assessed separately and together. In the pooled sample, a negative binomial mixed effects model was used, with family as a random variable to account for familial relatedness. The function used for probands and siblings alone was glm.nb() from the R package ‘MASS’ (58), and for pooled samples, glmmTMB() was used from the package ‘glmmTMB’. (59) All models used were adjusted for age, sex, ancestry, and in a second time for NVIQ and/or for the diagnosis of autism (Table S10).

See Table S7 for detail of the models used for the other phenotypical measures.

- *Computation of significance threshold*

To control for multiple testing, we computed our significance threshold by adapting the methodology of Cheverud (2001) previously applied to control the FWER when analyzing many SNPs (60). When computing the matrix of correlations between phenotypes, Cheverud argued that the eigenvalues of this matrix could be used to estimate the effective number of independent tests. Thus, we calculated an effective number of independent tests, *m_e_*, and then used this number in a Bonferroni-style correction.
*m_e_* was calculated as follow:

$$m_{e}=1+(m-1)\left( 1-\frac{1}{m} Var(\lambda) \right)$$

$$Var\left( \lambda\right)=\frac{1}{m-1} \sum_{i=1}^{m} {(\lambda_{i}-1)}^{2}$$

**SUPPLEMENTARY FIGURES**

**Figure S1: Distribution of NVIQ.**

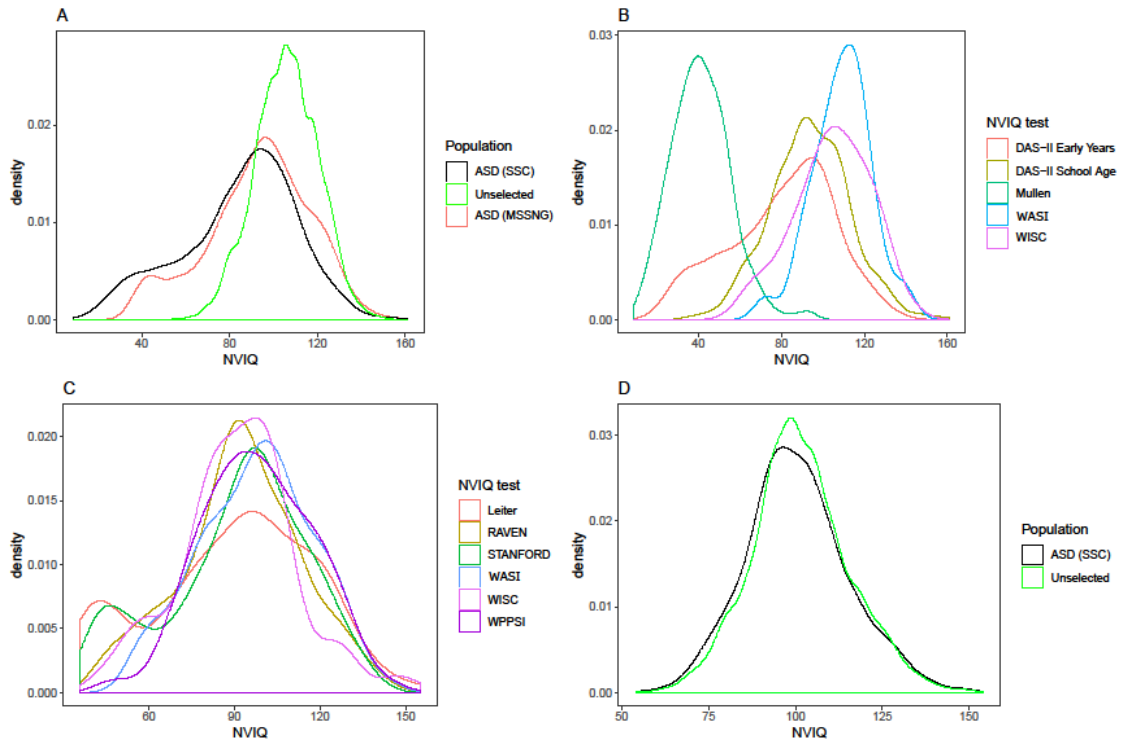
A) Density distribution of NVIQ in the autistic probands from SSC (black) and MSSNG (red), and from the unselected population (green: IMAGEN and SYS pooled). B) Density distribution of NVIQ in the autistic probands from SSC in function of the test used. C) Density distribution of NVIQ in the autistic probands from MSSNG in function of the test used. D) Density distribution of NVIQ in the autistic probands from SSC (black) and from the unselected population (green: IMAGEN and SYS pooled) after 1:1 matching procedure. NVIQ: Non-verbal IQ; DAS-II: Differential Ability Scales, 2nd Edition; WASI: Wechsler Abbreviated Scale Intelligence, WISC: Wechsler Intelligence Scale for Children; WPPSI: Wechsler Preschool and Primary Scale of Intelligence.

**Figure S2: Distribution of NVIQ for individual below or above the median (98), using probands from SSC and MSSNG, and the unselected populations.**

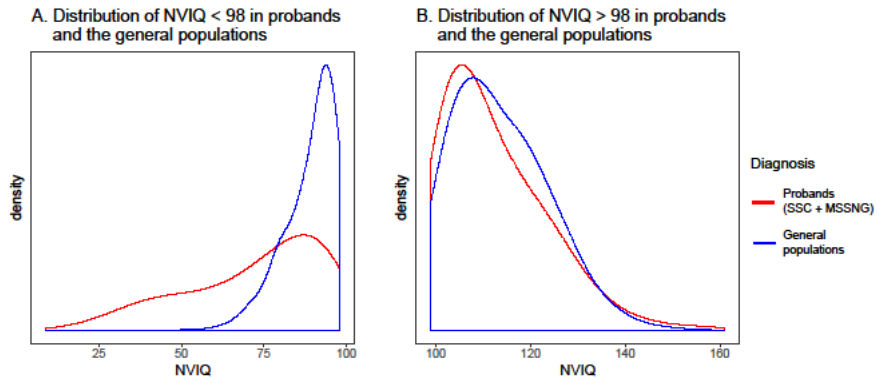

A) Density distribution of NVIQ in the autistic probands from SSC and MSSNG (red) and unselected populations (blue) with a NVIQ bellow 98. B) Density distribution of NVIQ in the autistic probands from SSC and MSSNG (red) and unselected populations (blue) with a NVIQ over 98.

**Figure S3: Effect of gene dosage measured by pLI on autism risk.**

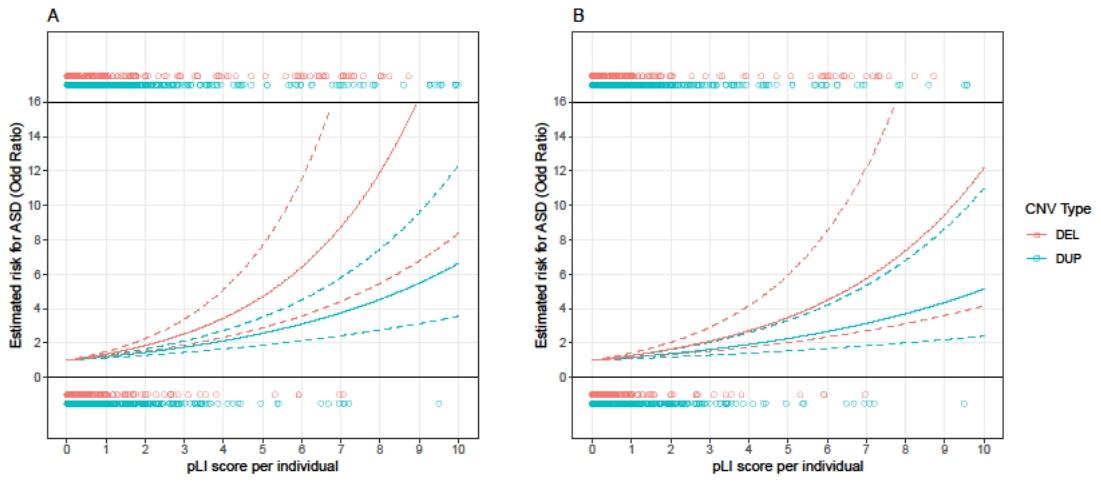

Representation of the non-linear effect of deleted (red) or duplicated (blue) point of pLI on autism risk estimated by logistic regression model when including (A) or excluding (B) the 16 recurrent CNVs detailed in Table S3. X-axis represents the sum of pLI score per individual and y-axis represents the risk of autism diagnosis estimated by our model in odds ratio.

**Figure S4: Effect of gene dosage measured by pLI on phenotypic measures adjusted for NVIQ in SSC probands.**

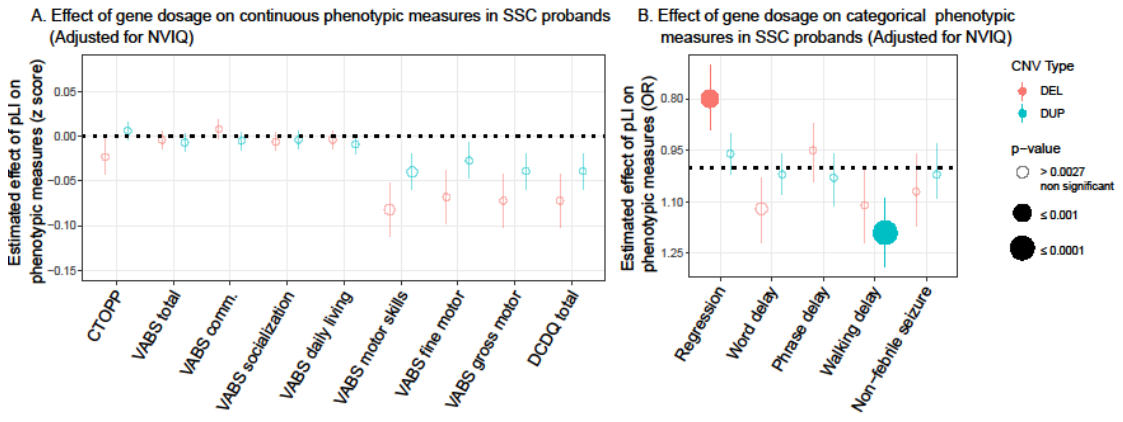

(A, B) Effect-size of a deleted (red) or duplicated (blue) point of pLI on continuous (A) and categorical (B) phenotypes in autistic probands from the SSC adjusted for NVIQ. Y-axis values of panel (A) are measures z-scored using normative data (Table S7) except for DCDQ which was z-scored using the SSC autistic proband group. Y-axis values of panel (B) are odds ratios computed by logistic regression. The significance threshold was computed (60) to account for multiple testing: 0.0027. CTOPP: Comprehensive Test of Phonological Processing; VABS-II: Vineland Adaptive behaviour Rating Scales - Second Edition; DCDQ: Developmental Coordination Disorder Questionnaire.

**Figure S5: Effect of gene dosage measured by pLI on age of onset for walking.**

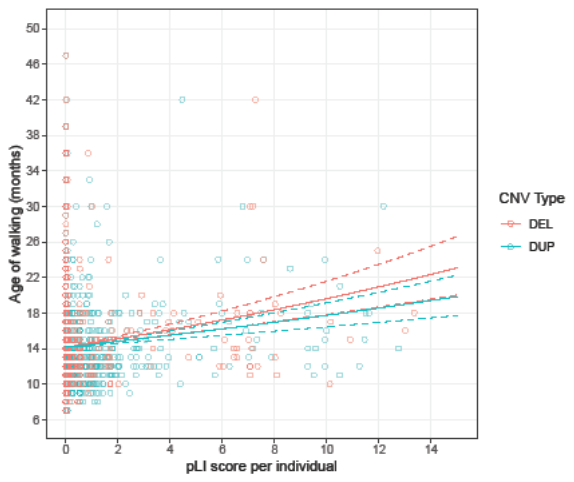

Representation of the non-linear effect of deleted (red: OR, 1.03 [95% CI, 1.02-1.04]; *P*=5×10^-12^) or duplicated (blue: OR, 1.02 [95% CI, 1.02-1.03]; *P*=2×10^-9^) point of pLI on age of onset for walking estimated using quasi-Poisson model adjusted for sex. X-axis represents the sum of pLI score per individual and y-axis represents the age of onset for walking in month.

**Figure S6: Effect of gene dosage measured by pLI on SRS score.**

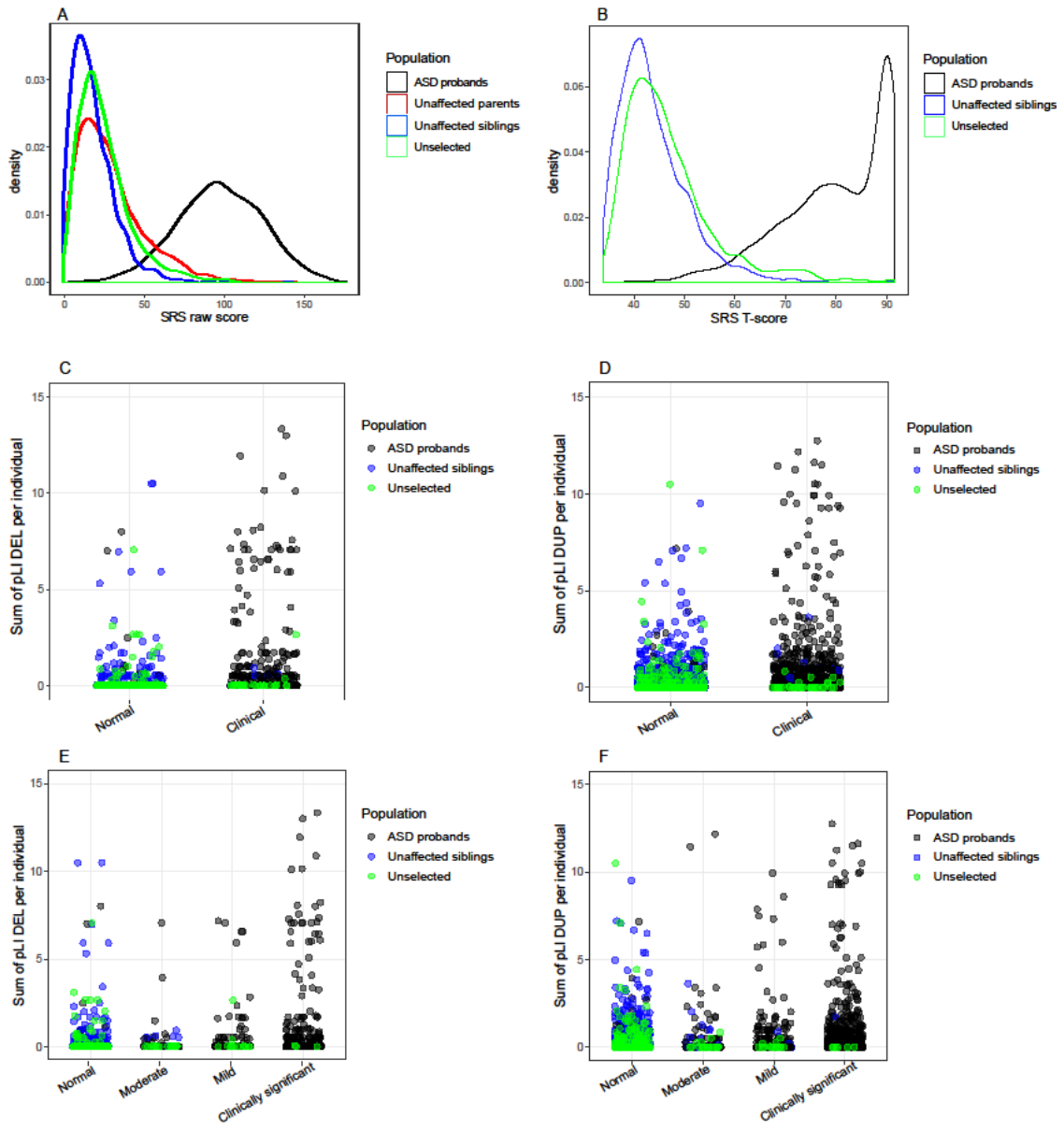

A) Density distribution of SRS raw scores in the autistic probands from SSC (black), their unaffected siblings (blue) and parents (red). B) Density distribution of SRS T-scores in the autistic probands from SSC (black), their unaffected siblings (blue), and from the unselected population (green: IMAGEN). C to F) Representation of the distribution of Sum of pLI deleted (C,E) or duplicated (D,F) per individual in function of SRS categories (binary (C,D) or all assessed categories (E,F)) in the autistic probands from SSC (black), unaffected siblings (blue) or unselected population (green: IMAGEN).

**SUPPLEMENTARY TABLES**

**Table S1: Description of cohorts for demographic, genotypical characteristics and phenotypical measures.**

| **Variables** | **IMAGEN** | **SYS** | **SSC probands** | | **SSC siblings** | | **SSC parents** | **MSSNG** |
| --- | --- | --- | --- | --- | --- | --- | --- | --- |
| Demographic characteristics | | | | | | | | |
| N individuals | 1,802 | 967 | 2,569 | 2,092 | | 5,138 | | 1,381 |
| Age, mean (SD)^(b)^ | 14.45 (0.37) | 14.99 (1.84) | 9.03 (3.58) | 10.06 (4.33) | | 41.48 (6.18) | | 9.21 (4.44) |
| N male (%) | 880 (48.91) | 462 (47.72) | 2,227 (86.66) | 969 (46.31) | | 2,569 (50.00) | | 1,106 (80.09) |
| Genotypical characteristics (Chr 1-22) | | | | | | | | |
| Detection technology | Illumina 610Kq or 660Wq array | Illumina 610Kq or HumanOmniExpress12 array | Illumina 1Mv1, 1Mv3 or Omni 2.5 array | | | | | Illumina HiSeq, HiSeq 2,500 or HiSeqX sequencing |
| N carriers of CNVs | 855 | 610 | 1,835 | 1,490 | | 2,040 | | 932 |
| N carriers of documented CNVs^(a)^ | 26 | 22 | 77 | 33 | | 82 | | 40 |
| N total CNVs | 2,712 | 2,213 | 6,597 | 5,159 | | 12,689 | | 4,075 |
| N deletions (total) | 1,574 | 1,212 | 3,323 | 2,562 | | 6,410 | | 2,435 |
| N deletions,  mean (SD) | 0.87 (0.91) | 1.25 (1.06) | 1.29 (1.05) | 1.22 (1.05) | | 1.24 (1.04) | | 1.76 (1.39) |
| N duplications (total) | 1,138 | 1,001 | 3,274 | 2,597 | | 6,279 | | 1,640 |
| N duplications,  mean (SD) | 0.63 (0.79) | 1.03 (1.04) | 1.27 (1.13) | 1.24 (1.12) | | 1.22 (1.11) | | 1.19 (1,15) |
| pLI deletions, sum | 94.97 | 55.16 | 416.00 | 118.08 | | 232.37 | | 171.61 |
| pLI deletions per individual, mean (SD) | 0.05 (0.34) | 0.06 (0.46) | 0.16 (1.00) | 0.06 (0.46) | | 0.04 (0.39) | | 0.12 (0.63) |
| pLI duplications, sum | 226.27 | 248.75 | 816.38 | 375.74 | | 891.78 | | 350.23 |
| pLI duplications per individual, mean (SD) | 0.13 (0.53) | 0.26 (0.73) | 0.32 (1.44) | 0.18 (0.63) | | 0.17 (0.60) | | 0.25 (1.12) |
| Phenotypical measures | | | | | | | | |
| N NVIQ | 1,744 | 966 | 2,569 | - | | - | | 1,381 |
| NVIQ,  mean (SD) | 106.62 (14.77) | 104.50 (13.09) | 84.47 (26.27) | - | | - | | 92.97 (23.72) |
| *Autism related symptoms* | | | | | | | | |
| N DAWBA DSM | 1,303 | - | - | - | | - | | - |
| N SRS | 977 | - | 2,556 | 2,078 | | 4,838 | | 598 |
| SRS raw score,  mean (SD) | 25.42 (16.99) | - | 97.98 (26.97) | 18.36 (13.82) | | 29.53 (21.34) | | - |
| SRS T-score,  mean (SD) | 49.86 (36.51) | - | 79.52 (10.45) | 43.80 (7.04) | | - | | - |
| N ADOS-overall css | - | - | 2,498 | - | | - | | 679 |
| ADOS-overall css,  mean (SD) | - | - | 7.45 (1.68) | - | | - | | 7.30 (2.11) |
| N ADOS-social affect css | - | - | 2,372 | - | | - | | 325 |
| ADOS-social css,  mean (SD) | - | - | 7.25 (1.76) | - | | - | | 6.86 (2.00) |
| N ADOS-rrsb css | - | - | 2,449 | - | | - | | 330 |
| ADOS-rrsb css,  mean (SD) | - | - | 7.78 (1.92) | - | | - | | 8.52 (1.51) |
| N ADIR reciprocal social interactions | - | - | 2,567 | - | | - | | 403 |
| ADIR reciprocal social interactions, mean (SD) | - | - | 20.35 (5.69) | - | | - | | 18.26 (7.98) |
| N ADIR rrb | - | - | 2,567 | - | | - | | 696 |
| ADIR rrb, mean (SD) | - | - | 6.52 (2.50) | - | | - | | 6.43 (2.50) |
| N ADIR verbal communication | - | - | 2,254 | - | | - | | 374 |
| ADIR verbal communication,  mean (SD) | - | - | 16.48 (4.29) | - | | - | | 14.42 (5.82) |
| N ADIR non-verbal communication | - | - | 2,567 | - | | - | | 71 |
| ADIR non-verbal communication,  mean (SD) | - | - | 9.26 (3.45) | - | | - | | 10.76 (3.03) |
| N with/without regression | - | - | 911/1,657 | - | | - | | - |
| *Language and phonology* | | | | | | | | |
| N ADIR overall level of language | - | - | 2,568 | - | | - | | 1,039 |
| N age of first word | - | - | 2,467 | - | | - | | - |
| age of first word in months, mean (SD) | - | - | 24.38 (14.97) | - | | - | | - |
| N with/without word delay | - | - | 935/1,632 | - | | - | | - |
| N age of first phrase | - | - | 2,311 | - | | - | | - |
| age of first phrase in months, mean (SD) | - | - | 39.18 (18.52) | - | | - | | - |
| N with/without phrase delay | - | - | 1,567/1,000 | - | | - | | - |
| N CTOPP | - | - | 1,988 | - | | - | | - |
| CTOPP, mean (SD) | - | - | 7.76 (2.86) | - | | - | | - |
| *behavioural problems* | | | | | | | | |
| N CBCL total score | - | - | 1,945 | 1,596 | | - | | - |
| CBCL total score,  mean (SD) | - | - | 50.77 (23.93) | 17.99 (15.18) | | - | | - |
| N CBCL internalizing | - | - | 1,945 | 1,596 | | - | | - |
| CBCL internalizing,  mean (SD) | - | - | 11.06 (8.46) | 4.71 (5.25) | | - | | - |
| N CBCL externalizing | - | - | 1,945 | 1,596 | | - | | - |
| CBCL externalizing,  mean (SD) | - | - | 11.56 (7.91) | 5.06 (5.05) | | - | | - |
| *Adaptative skills* | | | | | | | | |
| N VABS-II total | - | - | 2,569 | - | | - | | - |
| VABS-II total,  mean (SD) | - | - | 73.08 (12.13) | - | | - | | - |
| N VABS-II daily living | - | - | 2,569 | - | | - | | - |
| VABS-II daily living,  mean (SD) | - | - | 76.33 (13.92) | - | | - | | - |
| N VABS-II communication | - | - | 2,569 | - | | - | | - |
| VABS-II communication,  mean (SD) | - | - | 76.98 (14.63) | - | | - | | - |
| N VABS-II socialization | - | - | 2,569 | - | | - | | - |
| VABS-II socialization,  mean (SD) | - | - | 70.91 (12.59) | - | | - | | - |
| *Motor skills* | | | | | | | | |
| N VABS-II motor | - | - | 919 | - | | - | | - |
| VABS-II motor,  mean (SD) | - | - | 81.75 (12.60) | - | | - | | - |
| N VABS-II gross motor | - | - | 926 | - | | - | | - |
| VABS-II gross motor,  mean (SD) | - | - | 12.25 (2.23) | - | | - | | - |
| N VABS-II fine motor | - | - | 923 | - | | - | | - |
| VABS-II fine motor,  mean (SD) | - | - | 11.78 (2.69) | - | | - | | - |
| N age of onset of walking | - | - | 2,550 | - | | - | | - |
| age of onset of walking in months, mean (SD) | - | - | 13.56 (4.00) | - | | - | | - |
| N with/without onset of walking delay | - | - | 159/2,405 | - | | - | | - |
| N DCDQ | - | - | 2,209 | - | | - | | - |
| DCDQ, mean (SD) | - | - | 38.50 (12.44) | - | | - | | - |
| *Associated neurological condition* | | | | | | | | |
| N with/without non-febrile seizure | - | - | 233/2,333 | - | | - | | - |

(a) Number of carriers of a CNV which overlap ≥ 30% with a documented CNV from the Table S6; SD: Standard deviation; Sum pLI deletions or duplications: sum of score of pLI for the entire population; NVIQ: Non-verbal intelligence quotient; DAWBA: Development and Well-Being Assessment; SRS: Social Responsiveness Scale; ADOS: Autism Diagnostic Observation Schedule; css: calibrated severity score; rrsb: repetitive, restricted and stereotyped behaviours; ADI-R: Autism Diagnostic Interview-Revised; CTOPP: Comprehensive Test of Phonological Processing; CBCL: Child behaviour Checklist; VABS-II: Vineland Adaptive behaviour Rating Scales - Second Edition; DCDQ: Developmental Coordination Disorder Questionnaire.

**Table S2: NVIQ available in autistic probands from SSC and MSSNG, and individuals from unselected population.**

| **Cohorts** | **N available NVIQ** | **Age** | | **Males** | | **NVIQ** | | |
| --- | --- | --- | --- | --- | --- | --- | --- | --- |
|  |  | mean | SD | N | % | Test used | Mean | SD |
| IMAGEN | 1,744 | 14.45 | 0.37 | 880 | 48.91 | WISC-IV | 106.62 | 14.77 |
| SYS | 966 | 14.99 | 1.84 | 462 | 47.72 | WISC-III | 104.50 | 13.09 |
| SSC autistic probands | 2,564 | 9.03 | 3.58 | 2,227 | 86.66 | DAS-II, MSEL, WASI-I, WISC-IV | 84.47 | 26.27 |
| MSSNG autistic probands | 1,381 | 9.21 | 4.44 | 1,106 | 80.09 | Leiter, Raven, Stanford-Binet, WASI-I, WASI-II, WISC-IV, WPPSI-IV | 92.97 | 23.72 |

SD: Standard deviation; NVIQ: Non-verbal intelligence quotient; DAS-II: Differential Ability Scales - Second Edition; MSEL: Mullen Scale of Early Learning; WASI-I or II: Wechsler Abbreviated Scale of Intelligence – First or Second Edition; WISC-IV: Wechsler Intelligence Scale for Children, Fourth Edition; WPPSI-IV: Wechsler Preschool and Primary Scale of Intelligence – Fourth Edition.

**Table S3: Breakpoints used to detect recurrent CNVs associated to neurodevelopmental disorders.**

| **Reference** | **Chr** | **Start hg19** | **Stop hg19** | **Type** | **Note protective CNV** |
| --- | --- | --- | --- | --- | --- |
| Cooper et al. 2012 (61) and Coe et al. 2014 (62) | chr1 | 1 | 10077413 | DEL | *GABRD* |
| Cooper et al. 2012 (61) and Coe et al. 2014 (62) | chr1 | 710137 | 9977413 | DEL |  |
| Cooper et al. 2012 (61) and Coe et al. 2014 (62) | chr1 | 860137 | 3660140 | DUP |  |
| Cooper et al. 2012 (61) and Coe et al. 2014 (62) | chr1 | 145288643 | 145628643 | DEL | *HFE2* |
| Cooper et al. 2012 (61) and Coe et al. 2014 (62) | chr1 | 145338643 | 149783376 | DUP |  |
| Cooper et al. 2012 (61) and Coe et al. 2014 (62) | chr1 | 145338643 | 147883376 | DEL |  |
| Cooper et al. 2012 (61) and Coe et al. 2014 (62) | chr1 | 146573376 | 147393376 | DEL | *GJA5* |
| Cooper et al. 2012 (61) and Coe et al. 2014 (62) | chr1 | 146573376 | 147393376 | DUP |  |
| Cooper et al. 2012 (61) and Coe et al. 2014 (62) | chr1 | 168733376 | 173733377 | DEL | *FMO* and *DNM3* |
| Cooper et al. 2012 (61) and Coe et al. 2014 (62) | chr1 | 245033377 | 248833377 | DUP |  |
| Cooper et al. 2012 (61) and Coe et al. 2014 (62) | chr2 | 50146496 | 51256496 | DEL | *NRXN1* |
| Cooper et al. 2012 (61) and Coe et al. 2014 (62) | chr2 | 57746496 | 61736496 | DEL | *VRK2* |
| Cooper et al. 2012 (61) and Coe et al. 2014 (62) | chr2 | 59646496 | 63146496 | DUP | *PEX13* to *AHSA2* |
| Cooper et al. 2012 (61) and Coe et al. 2014 (62) | chr2 | 96726273 | 97676273 | DUP |  |
| Cooper et al. 2012 (61) and Coe et al. 2014 (62) | chr2 | 100693568 | 108443568 | DEL | *NCK2* and *FHL2* |
| Cooper et al. 2012 (61) and Coe et al. 2014 (62) | chr2 | 111333937 | 113233529 | DEL |  |
| Cooper et al. 2012 (61) and Coe et al. 2014 (62) | chr2 | 111383531 | 113093529 | DEL |  |
| Cooper et al. 2012 (61) and Coe et al. 2014 (62) | chr2 | 111383531 | 113093529 | DUP |  |
| Cooper et al. 2012 (61) and Coe et al. 2014 (62) | chr2 | 200161755 | 200511755 | DEL | *SATB2* |
| Cooper et al. 2012 (61) and Coe et al. 2014 (62) | chr2 | 235735261 | 243102476 | DEL |  |
| Cooper et al. 2012 (61) and Coe et al. 2014 (62) | chr2 | 239705243 | 242471327 | DEL | *HDAC4* |
| Cooper et al. 2012 (61) and Coe et al. 2014 (62) | chr3 | 9525000 | 11025000 | DUP | *JAGN1* to *TATDN2* |
| Cooper et al. 2012 (61) and Coe et al. 2014 (62) | chr3 | 87237310 | 87557310 | DEL | *CHMP2B* to *POU1F1* |
| Cooper et al. 2012 (61) and Coe et al. 2014 (62) | chr3 | 115237310 | 115647310 | DEL | *GAP43* |
| Cooper et al. 2012 (61) and Coe et al. 2014 (62) | chr3 | 191517306 | 193017306 | DEL | *FGF12* |
| Cooper et al. 2012 (61) and Coe et al. 2014 (62) | chr3 | 195715603 | 197355603 | DEL |  |
| Cooper et al. 2012 (61) and Coe et al. 2014 (62) | chr3 | 195715603 | 197355603 | DUP |  |
| Cooper et al. 2012 (61) and Coe et al. 2014 (62) | chr3 | 195745603 | 197355603 | DEL | *DLG1* |
| Cooper et al. 2012 (61) and Coe et al. 2014 (62) | chr3 | 195745603 | 197355603 | DUP |  |
| Cooper et al. 2012 (61) and Coe et al. 2014 (62) | chr4 | 110000 | 7049099 | DEL |  |
| Cooper et al. 2012 (61) and Coe et al. 2014 (62) | chr4 | 1870202 | 2010202 | DEL | *WHSC1* and *WHSC2* |
| Cooper et al. 2012 (61) and Coe et al. 2014 (62) | chr4 | 80780976 | 83280976 | DEL |  |
| Cooper et al. 2012 (61) and Coe et al. 2014 (62) | chr5 | 1 | 11727000 | DEL |  |
| Cooper et al. 2012 (61) and Coe et al. 2014 (62) | chr5 | 87964244 | 88224244 | DEL | *MEF2C* |
| Cooper et al. 2012 (61) and Coe et al. 2014 (62) | chr5 | 175717394 | 177057394 | DEL | *NSD1* |
| Cooper et al. 2012 (61) and Coe et al. 2014 (62) | chr5 | 180117394 | 180817394 | DEL |  |
| Cooper et al. 2012 (61) and Coe et al. 2014 (62) | chr6 | 92043279 | 104693307 | DEL | *FOXP1* and *SIM1* |
| Cooper et al. 2012 (61) and Coe et al. 2014 (62) | chr6 | 100813279 | 100943279 | DEL |  |
| Cooper et al. 2012 (61) and Coe et al. 2014 (62) | chr6 | 165330010 | 170908075 | DEL |  |
| Cooper et al. 2012 (61) and Coe et al. 2014 (62) | chr7 | 10239 | 3833474 | DUP |  |
| Cooper et al. 2012 (61) and Coe et al. 2014 (62) | chr7 | 66482565 | 72272064 | DEL | *AUTS2* |
| Cooper et al. 2012 (61) and Coe et al. 2014 (62) | chr7 | 66482565 | 72272064 | DUP |  |
| Cooper et al. 2012 (61) and Coe et al. 2014 (62) | chr7 | 72662064 | 74262064 | DUP |  |

**Table S3 continued**

| **Reference** | **Chr** | **Start hg19** | **Stop hg19** | **Type** | **Note protective CNV** |
| --- | --- | --- | --- | --- | --- |
| Cooper et al. 2012 (61) and Coe et al. 2014 (62) | chr7 | 72662064 | 74262064 | DEL |  |
| Cooper et al. 2012 (61) and Coe et al. 2014 (62) | chr7 | 72742064 | 74142064 | DUP |  |
| Cooper et al. 2012 (61) and Coe et al. 2014 (62) | chr7 | 72742064 | 74142064 | DEL | *ELN* and *GTF2I* |
| Cooper et al. 2012 (61) and Coe et al. 2014 (62) | chr7 | 74962064 | 76662064 | DEL | *RHBDD2* and *HIP1* |
| Cooper et al. 2012 (61) and Coe et al. 2014 (62) | chr7 | 74962064 | 76662064 | DUP |  |
| Cooper et al. 2012 (61) and Coe et al. 2014 (62) | chr8 | 160000 | 11912591 | DEL |  |
| Cooper et al. 2012 (61) and Coe et al. 2014 (62) | chr8 | 8092590 | 11892591 | DEL | *SOX7* and *CLDN23* |
| Cooper et al. 2012 (61) and Coe et al. 2014 (62) | chr8 | 8092590 | 11892591 | DUP |  |
| Cooper et al. 2012 (61) and Coe et al. 2014 (62) | chr8 | 8212590 | 11912591 | DUP |  |
| Cooper et al. 2012 (61) and Coe et al. 2014 (62) | chr9 | 32010000 | 39010000 | DUP |  |
| Cooper et al. 2012 (61) and Coe et al. 2014 (62) | chr9 | 131060179 | 141080179 | DEL |  |
| Cooper et al. 2012 (61) and Coe et al. 2014 (62) | chr9 | 137810179 | 141080179 | DEL | *EHMT1* |
| Cooper et al. 2012 (61) and Coe et al. 2014 (62) | chr9 | 137810179 | 141080179 | DUP |  |
| Cooper et al. 2012 (61) and Coe et al. 2014 (62) | chr9 | 137860179 | 141080179 | DEL |  |
| Cooper et al. 2012 (61) and Coe et al. 2014 (62) | chr9 | 137860179 | 141080179 | DUP |  |
| Cooper et al. 2012 (61) and Coe et al. 2014 (62) | chr10 | 46929994 | 48429994 | DEL |  |
| Cooper et al. 2012 (61) and Coe et al. 2014 (62) | chr10 | 49389994 | 52389994 | DUP |  |
| Cooper et al. 2012 (61) and Coe et al. 2014 (62) | chr10 | 81690020 | 88940020 | DEL | *SFTPD* to *GLUD1*, *NRG3* |
| Cooper et al. 2012 (61) and Coe et al. 2014 (62) | chr10 | 81960020 | 88800020 | DEL | *NRG3* and *GRID1* |
| Cooper et al. 2012 (61) and Coe et al. 2014 (62) | chr10 | 127760010 | 135400010 | DEL |  |
| Cooper et al. 2012 (61) and Coe et al. 2014 (62) | chr11 | 310000 | 3443424 | DEL |  |
| Cooper et al. 2012 (61) and Coe et al. 2014 (62) | chr11 | 43983424 | 46063424 | DEL | *EXT2* |
| Cooper et al. 2012 (61) and Coe et al. 2014 (62) | chr11 | 67753424 | 71282352 | DEL | *SHANK2* |
| Cooper et al. 2012 (61) and Coe et al. 2014 (62) | chr11 | 128044790 | 134844790 | DEL |  |
| Cooper et al. 2012 (61) and Coe et al. 2014 (62) | chr12 | 6469739 | 6809739 | DUP | *SCNN1A* to *PIANP* |
| Cooper et al. 2012 (61) and Coe et al. 2014 (62) | chr12 | 65073733 | 68643733 | DEL | *GRIP1* and *HMGA2* |
| Cooper et al. 2012 (61) and Coe et al. 2014 (62) | chr14 | 104480247 | 106378955 | DEL |  |
| Cooper et al. 2012 (61) and Coe et al. 2014 (62) | chr15 | 22648636 | 28626405 | DEL |  |
| Cooper et al. 2012 (61) and Coe et al. 2014 (62) | chr15 | 22648636 | 31912708 | DUP |  |
| Cooper et al. 2012 (61) and Coe et al. 2014 (62) | chr15 | 22798636 | 23088559 | DEL | *NIPA1* |
| Cooper et al. 2012 (61) and Coe et al. 2014 (62) | chr15 | 24818907 | 28426405 | DEL |  |
| Cooper et al. 2012 (61) and Coe et al. 2014 (62) | chr15 | 24818907 | 28426405 | DUP |  |
| Cooper et al. 2012 (61) and Coe et al. 2014 (62) | chr15 | 30862708 | 32962708 | DEL |  |
| Cooper et al. 2012 (61) and Coe et al. 2014 (62) | chr15 | 31132708 | 32482708 | DEL | *CHRNA7* |
| Cooper et al. 2012 (61) and Coe et al. 2014 (62) | chr15 | 72912946 | 75792945 | DEL | *PML* |
| Cooper et al. 2012 (61) and Coe et al. 2014 (62) | chr15 | 72912946 | 74412947 | DEL | *BBS4, NPTN, NEO1* |
| Cooper et al. 2012 (61) and Coe et al. 2014 (62) | chr15 | 72962947 | 75532947 | DEL |  |
| Cooper et al. 2012 (61) and Coe et al. 2014 (62) | chr15 | 72962947 | 76012945 | DEL |  |
| Cooper et al. 2012 (61) and Coe et al. 2014 (62) | chr15 | 74012947 | 75532947 | DEL |  |
| Cooper et al. 2012 (61) and Coe et al. 2014 (62) | chr15 | 74012947 | 76012945 | DEL |  |
| Cooper et al. 2012 (61) and Coe et al. 2014 (62) | chr15 | 74012947 | 78132945 | DEL |  |
| Cooper et al. 2012 (61) and Coe et al. 2014 (62) | chr15 | 74012947 | 75532947 | DEL |  |
| Cooper et al. 2012 (61) and Coe et al. 2014 (62) | chr15 | 74412947 | 75592947 | DEL | *CLK3, CSK* |
| Cooper et al. 2012 (61) and Coe et al. 2014 (62) | chr15 | 75592947 | 75792945 | DEL | *SIN3A* |

**Table S3 continued**

| **Reference** | **Chr** | **Start hg19** | **Stop hg19** | **Type** | **Note protective CNV** |
| --- | --- | --- | --- | --- | --- |
| Cooper et al. 2012 (61) and Coe et al. 2014 (62) | chr15 | 75972945 | 78202945 | DEL | *FBXO22* and *TPSAN3* |
| Cooper et al. 2012 (61) and Coe et al. 2014 (62) | chr15 | 83182945 | 84738996 | DEL | *HOMER2* and *BNC1* |
| Cooper et al. 2012 (61) and Coe et al. 2014 (62) | chr15 | 85138996 | 85698996 | DEL |  |
| Cooper et al. 2012 (61) and Coe et al. 2014 (62) | chr15 | 99362477 | 102521392 | DEL |  |
| Cooper et al. 2012 (61) and Coe et al. 2014 (62) | chr15 | 99362477 | 102521392 | DUP |  |
| Cooper et al. 2012 (61) and Coe et al. 2014 (62) | chr16 | 160000 | 5159999 | DEL |  |
| Cooper et al. 2012 (61) and Coe et al. 2014 (62) | chr16 | 3779999 | 3859999 | DEL |  |
| Cooper et al. 2012 (61) and Coe et al. 2014 (62) | chr16 | 14892499 | 16892499 | DUP |  |
| Cooper et al. 2012 (61) and Coe et al. 2014 (62) | chr16 | 14892499 | 18292499 | DEL |  |
| Cooper et al. 2012 (61) and Coe et al. 2014 (62) | chr16 | 15502499 | 16292499 | DEL | *MYH11* |
| Cooper et al. 2012 (61) and Coe et al. 2014 (62) | chr16 | 15502499 | 16292499 | DUP |  |
| Cooper et al. 2012 (61) and Coe et al. 2014 (62) | chr16 | 21352499 | 29442499 | DEL |  |
| Cooper et al. 2012 (61) and Coe et al. 2014 (62) | chr16 | 21612499 | 29042499 | DEL |  |
| Cooper et al. 2012 (61) and Coe et al. 2014 (62) | chr16 | 21892499 | 22492499 | DEL |  |
| Cooper et al. 2012 (61) and Coe et al. 2014 (62) | chr16 | 21942499 | 22462499 | DEL | *EEF2K* and *CDR2* |
| Cooper et al. 2012 (61) and Coe et al. 2014 (62) | chr16 | 28442499 | 30342499 | DUP |  |
| Cooper et al. 2012 (61) and Coe et al. 2014 (62) | chr16 | 28442499 | 30342499 | DEL |  |
| Cooper et al. 2012 (61) and Coe et al. 2014 (62) | chr16 | 28772499 | 29112499 | DEL | *SH2B1* |
| Cooper et al. 2012 (61) and Coe et al. 2014 (62) | chr16 | 29652499 | 30202499 | DEL | *TBX6* |
| Cooper et al. 2012 (61) and Coe et al. 2014 (62) | chr16 | 29652499 | 30202499 | DUP |  |
| Cooper et al. 2012 (61) and Coe et al. 2014 (62) | chr16 | 83792499 | 90222499 | DEL |  |
| Cooper et al. 2012 (61) and Coe et al. 2014 (62) | chr17 | 50000 | 2593250 | DEL | *YWHAE* and *PAFAH1B1* |
| Cooper et al. 2012 (61) and Coe et al. 2014 (62) | chr17 | 50000 | 2593250 | DUP | *YWHAE* and *PAFAH1B1* |
| Cooper et al. 2012 (61) and Coe et al. 2014 (62) | chr17 | 100000 | 4153251 | DEL |  |
| Cooper et al. 2012 (61) and Coe et al. 2014 (62) | chr17 | 553250 | 1353250 | DEL | *PAFAH1B1* |
| Cooper et al. 2012 (61) and Coe et al. 2014 (62) | chr17 | 553250 | 1353250 | DUP | *PAFAH1B1* |
| Cooper et al. 2012 (61) and Coe et al. 2014 (62) | chr17 | 2363250 | 2923250 | DEL | *YWHAE* |
| Cooper et al. 2012 (61) and Coe et al. 2014 (62) | chr17 | 2363250 | 2923250 | DUP | *YWHAE* |
| Cooper et al. 2012 (61) and Coe et al. 2014 (62) | chr17 | 16709275 | 20479408 | DUP |  |
| Cooper et al. 2012 (61) and Coe et al. 2014 (62) | chr17 | 16709275 | 20309408 | DEL |  |
| Cooper et al. 2012 (61) and Coe et al. 2014 (62) | chr17 | 16709275 | 20479408 | DEL |  |
| Cooper et al. 2012 (61) and Coe et al. 2014 (62) | chr17 | 29165874 | 30215887 | DEL | *NF1* |
| Cooper et al. 2012 (61) and Coe et al. 2014 (62) | chr17 | 34815887 | 36205887 | DUP |  |
| Cooper et al. 2012 (61) and Coe et al. 2014 (62) | chr17 | 34815887 | 36205887 | DEL | *TCF2* |
| Cooper et al. 2012 (61) and Coe et al. 2014 (62) | chr17 | 43644217 | 44144178 | DEL |  |
| Cooper et al. 2012 (61) and Coe et al. 2014 (62) | chr17 | 43704217 | 44184217 | DUP |  |
| Cooper et al. 2012 (61) and Coe et al. 2014 (62) | chr17 | 43704217 | 44184217 | DEL | *MAPT* |
| Cooper et al. 2012 (61) and Coe et al. 2014 (62) | chr17 | 57655218 | 58075218 | DEL | *TUBD1* and *TMEM49* |
| Cooper et al. 2012 (61) and Coe et al. 2014 (62) | chr17 | 58065218 | 60305218 | DEL | *TBX2* and *TBX4* |
| Cooper et al. 2012 (61) and Coe et al. 2014 (62) | chr17 | 72088405 | 81060000 | DEL |  |
| Cooper et al. 2012 (61) and Coe et al. 2014 (62) | chr18 | 110000 | 5310000 | DEL |  |
| Cooper et al. 2012 (61) and Coe et al. 2014 (62) | chr18 | 70949020 | 77899009 | DEL |  |
| Cooper et al. 2012 (61) and Coe et al. 2014 (62) | chr19 | 199000 | 5899000 | DUP |  |

**Table S3 continued**

| **Reference** | **Chr** | **Start hg19** | **Stop hg19** | **Type** | **Note protective CNV** |
| --- | --- | --- | --- | --- | --- |
| Cooper et al. 2012 (61) and Coe et al. 2014 (62) | chr19 | 199000 | 8789000 | DEL |  |
| Cooper et al. 2012 (61) and Coe et al. 2014 (62) | chr21 | 42478130 | 47975572 | DEL |  |
| Cooper et al. 2012 (61) and Coe et al. 2014 (62) | chr22 | 17470000 | 25020000 | DUP |  |
| Cooper et al. 2012 (61) and Coe et al. 2014 (62) | chr22 | 18820000 | 22270000 | DEL |  |
| Cooper et al. 2012 (61) and Coe et al. 2014 (62) | chr22 | 19020000 | 20290000 | DUP |  |
| Cooper et al. 2012 (61) and Coe et al. 2014 (62) | chr22 | 19020000 | 20290000 | DEL |  |
| Cooper et al. 2012 (61) and Coe et al. 2014 (62) | chr22 | 21910000 | 23650000 | DEL | *BCR* and *MAPK1* |
| Cooper et al. 2012 (61) and Coe et al. 2014 (62) | chr22 | 21910000 | 23650000 | DUP |  |
| Cooper et al. 2012 (61) and Coe et al. 2014 (62) | chr22 | 44268667 | 51244566 | DEL |  |
| Cooper et al. 2012 (61) and Coe et al. 2014 (62) | chr22 | 47021336 | 51244566 | DUP |  |
| Cooper et al. 2012 (61) and Coe et al. 2014 (62) | chr22 | 51113134 | 51173134 | DEL | *SHANK3* |
| Marshall et al 2017 (63) | chr1 | 145430996 | 148237104 | DEL |  |
| Marshall et al 2017 (63) | chr1 | 145430996 | 148237104 | DUP |  |
| Marshall et al 2017 (63) | chr2 | 49920350 | 51032536 | DEL | *NRXN1* |
| Marshall et al 2017 (63) | chr3 | 196018732 | 197628732 | DEL |  |
| Marshall et al 2017 (63) | chr7 | 65373855 | 65401085 | DEL | *ZNF92,* Protective |
| Marshall et al 2017 (63) | chr7 | 65373855 | 65401085 | DUP | *ZNF92,* Protective |
| Marshall et al 2017 (63) | chr7 | 73328061 | 74727726 | DUP |  |
| Marshall et al 2017 (63) | chr7 | 158660506 | 159179546 | DEL | *WDR60* and *VIPR2* |
| Marshall et al 2017 (63) | chr7 | 158660506 | 159179546 | DUP | *WDR60* and *VIPR2* |
| Marshall et al 2017 (63) | chr8 | 100025494 | 100889814 | DEL | *VPS13B* |
| Marshall et al 2017 (63) | chr9 | 841690 | 969090 | DEL | *DMRT1* |
| Marshall et al 2017 (63) | chr9 | 841690 | 969090 | DUP | *DMRT1* |
| Marshall et al 2017 (63) | chr13 | 20397624 | 20437776 | DUP | *ZMYM5,* Protective |
| Marshall et al 2017 (63) | chr15 | 22784509 | 23074432 | DEL |  |
| Marshall et al 2017 (63) | chr15 | 30840505 | 32190507 | DEL |  |
| Marshall et al 2017 (63) | chr16 | 28811178 | 29041178 | DEL |  |
| Marshall et al 2017 (63) | chr16 | 29641178 | 30191178 | DUP |  |
| Marshall et al 2017 (63) | chr22 | 19032487 | 21065711 | DEL |  |
| Marshall et al 2017 (63) | chr22 | 19032487 | 21065711 | DUP | Protective |
| Marshall et al 2017 (63) | chrX | 148793685 | 148798928 | DUP | *MAGEA11*, Protective |
| Marshall et al 2017 (63) | chrX | 154918531 | 155342497 | DUP |  |
| Moreno del luca et al 2013 (64) | chr1 | 144000000 | 144340000 | DUP |  |
| Moreno del luca et al 2013 (64) | chr1 | 144000000 | 144340000 | DEL |  |
| Moreno del luca et al 2013 (64) | chr1 | 145040000 | 145860000 | DEL |  |
| Moreno del luca et al 2013 (64) | chr1 | 145040000 | 145860000 | DUP |  |
| Moreno del luca et al 2013 (64) | chr1 | 145040000 | 145860000 | DEL |  |
| Moreno del luca et al 2013 (64) | chr1 | 145040000 | 145860000 | DUP |  |
| Moreno del luca et al 2013 (64) | chr3 | 197230000 | 198840000 | DUP |  |
| Moreno del luca et al 2013 (64) | chr3 | 197230000 | 198840000 | DEL |  |
| Moreno del luca et al 2013 (64) | chr5 | 175650000 | 176990000 | DUP |  |
| Moreno del luca et al 2013 (64) | chr5 | 175650000 | 176990000 | DEL |  |
| Moreno del luca et al 2013 (64) | chr7 | 72380000 | 73780000 | DEL |  |
| Moreno del luca et al 2013 (64) | chr7 | 72380000 | 73780000 | DUP |  |

**Table S3 continued**

| **Reference** | **Chr** | **Start hg19** | **Stop hg19** | **Type** | **Note protective CNV** |
| --- | --- | --- | --- | --- | --- |
| Moreno del luca et al 2013 (64) | chr8 | 8130000 | 11930000 | DEL |  |
| Moreno del luca et al 2013 (64) | chr8 | 8130000 | 11930000 | DUP |  |
| Moreno del luca et al 2013 (64) | chr10 | 81950000 | 88790000 | DUP |  |
| Moreno del luca et al 2013 (64) | chr15 | 22370000 | 26100000 | DEL |  |
| Moreno del luca et al 2013 (64) | chr15 | 22370000 | 26100000 | DUP |  |
| Moreno del luca et al 2013 (64) | chr15 | 22370000 | 26100000 | DUP |  |
| Moreno del luca et al 2013 (64) | chr15 | 28920000 | 30270000 | DEL |  |
| Moreno del luca et al 2013 (64) | chr15 | 28920000 | 30270000 | DUP |  |
| Moreno del luca et al 2013 (64) | chr15 | 28920000 | 30270000 | DEL |  |
| Moreno del luca et al 2013 (64) | chr15 | 28920000 | 30270000 | DUP |  |
| Moreno del luca et al 2013 (64) | chr16 | 15410000 | 16200000 | DUP |  |
| Moreno del luca et al 2013 (64) | chr16 | 15410000 | 16200000 | DEL |  |
| Moreno del luca et al 2013 (64) | chr16 | 21850000 | 22370000 | DUP |  |
| Moreno del luca et al 2013 (64) | chr16 | 21850000 | 22370000 | DEL |  |
| Moreno del luca et al 2013 (64) | chr16 | 28680000 | 29020000 | DEL |  |
| Moreno del luca et al 2013 (64) | chr16 | 28680000 | 29020000 | DUP |  |
| Moreno del luca et al 2013 (64) | chr16 | 28680000 | 29020000 | DEL |  |
| Moreno del luca et al 2013 (64) | chr16 | 28680000 | 29020000 | DUP |  |
| Moreno del luca et al 2013 (64) | chr16 | 29560000 | 30110000 | DEL |  |
| Moreno del luca et al 2013 (64) | chr16 | 29560000 | 30110000 | DUP |  |
| Moreno del luca et al 2013 (64) | chr16 | 29560000 | 30110000 | DEL |  |
| Moreno del luca et al 2013 (64) | chr16 | 29560000 | 30110000 | DUP |  |
| Moreno del luca et al 2013 (64) | chr17 | 16650000 | 20420000 | DEL |  |
| Moreno del luca et al 2013 (64) | chr17 | 16650000 | 20420000 | DUP |  |
| Moreno del luca et al 2013 (64) | chr17 | 26190000 | 27240000 | DUP |  |
| Moreno del luca et al 2013 (64) | chr17 | 31890000 | 33280000 | DEL |  |
| Moreno del luca et al 2013 (64) | chr17 | 31890000 | 33280000 | DUP |  |
| Moreno del luca et al 2013 (64) | chr17 | 31890000 | 33280000 | DEL |  |
| Moreno del luca et al 2013 (64) | chr17 | 41060000 | 41540000 | DUP |  |
| Moreno del luca et al 2013 (64) | chr22 | 17400000 | 18670000 | DEL |  |
| Moreno del luca et al 2013 (64) | chr22 | 17400000 | 18670000 | DUP |  |
| Moreno del luca et al 2013 (64) | chr22 | 17400000 | 18670000 | DEL |  |
| Moreno del luca et al 2013 (64) | chr22 | 17400000 | 18670000 | DUP |  |
| Moreno del luca et al 2013 (64) | chr22 | 20240000 | 21980000 | DEL |  |
| Moreno del luca et al 2013 (64) | chr22 | 20240000 | 21980000 | DUP |  |
| Stefansson et al. (65) | chr1 | 146089254 | 147859944 | DEL |  |
| Stefansson et al. (65) | chr1 | 146089254 | 147858944 | DUP |  |
| Stefansson et al. (65) | chr2 | 1792885 | 2335045 | DUP | *MYT1L* |
| Stefansson et al. (65) | chr2 | 50145643 | 51259674 | DEL | *NRXN1* |
| Stefansson et al. (65) | chr3 | 195766737 | 197216349 | DEL |  |
| Stefansson et al. (65) | chr7 | 157860945 | 159119486 | DEL | *WDR60* and *VIPR2* |
| Stefansson et al. (65) | chr7 | 158726462 | 158947294 | DUP | *WDR60* and *VIPR2* |
| Stefansson et al. (65) | chr10 | 46508694 | 51912781 | DEL |  |

**Table S3 continued**

| **Reference** | **Chr** | **Start hg19** | **Stop hg19** | **Type** | **Note protective CNV** |
| --- | --- | --- | --- | --- | --- |
| Stefansson et al. (65) | chr10 | 47543322 | 51912781 | DUP |  |
| Stefansson et al. (65) | chr13 | 93879078 | 95060273 | DUP | *GPC6* |
| Stefansson et al. (65) | chr15 | 22750305 | 23272733 | DEL |  |
| Stefansson et al. (65) | chr15 | 22770994 | 28535266 | DUP |  |
| Stefansson et al. (65) | chr15 | 28973396 | 30556183 | DUP |  |
| Stefansson et al. (65) | chr15 | 29562640 | 30689724 | DEL |  |
| Stefansson et al. (65) | chr15 | 30936285 | 32515849 | DEL |  |
| Stefansson et al. (65) | chr15 | 32018731 | 32620127 | DEL | *CHRNA7* |
| Stefansson et al. (65) | chr16 | 14989844 | 16291983 | DUP |  |
| Stefansson et al. (65) | chr16 | 15125441 | 16291983 | DEL |  |
| Stefansson et al. (65) | chr16 | 21947230 | 22423698 | DEL |  |
| Stefansson et al. (65) | chr16 | 28814098 | 29043450 | DEL |  |
| Stefansson et al. (65) | chr16 | 29595483 | 30192561 | DEL |  |
| Stefansson et al. (65) | chr16 | 29624247 | 30198151 | DUP |  |
| Stefansson et al. (65) | chr17 | 14101029 | 15471179 | DEL |  |
| Stefansson et al. (65) | chr17 | 34815551 | 36249430 | DEL |  |
| Stefansson et al. (65) | chr17 | 34815551 | 36249430 | DUP |  |
| Stefansson et al. (65) | chr22 | 20718116 | 21465780 | DUP |  |
| Stefansson et al. (65) | chr22 | 20733495 | 21465780 | DEL |  |
| Huguet et al. (3) | chr1 | 145430996 | 148237104 | DEL |  |
| Huguet et al. (3) | chr1 | 145430996 | 148237104 | DUP |  |
| Huguet et al. (3) | chr2 | 49920350 | 51032536 | DEL | *NRXN1* |
| Huguet et al. (3) | chr3 | 196018732 | 197628732 | DEL |  |
| Huguet et al. (3) | chr7 | 73328061 | 74727726 | DUP |  |
| Huguet et al. (3) | chr7 | 65373855 | 65401085 | DEL | *ZNF92* |
| Huguet et al. (3) | chr7 | 65373855 | 65401085 | DUP | *ZNF92* |
| Huguet et al. (3) | chr7 | 158660506 | 159179546 | DEL | *VIPR2* and *WDR60* |
| Huguet et al. (3) | chr7 | 158660506 | 159179546 | DUP | *VIPR2* and *WDR60* |
| Huguet et al. (3) | chr8 | 99013266 | 99877580 | DEL | *VPS13B* |
| Huguet et al. (3) | chr9 | 841690 | 969090 | DEL | *DMRT1* |
| Huguet et al. (3) | chr9 | 841690 | 969090 | DUP | *DMRT1* |
| Huguet et al. (3) | chr13 | 19837453 | 19863633 | DUP | *ZMYM5* |
| Huguet et al. (3) | chr15 | 30840505 | 32190507 | DEL |  |
| Huguet et al. (3) | chr15 | 22784509 | 23074432 | DEL |  |
| Huguet et al. (3) | chr16 | 29641178 | 30191178 | DUP |  |
| Huguet et al. (3) | chr16 | 28811178 | 29041178 | DEL |  |
| Huguet et al. (3) | chr22 | 19032487 | 21065711 | DEL |  |
| Huguet et al. (3) | chr22 | 19032487 | 21065711 | DUP |  |

Chr: Chromosome; hg19: Homo sapiens (human) genome assembly GRCh37 (hg19) from Genome Reference Consortium; DEL: deletion; DUP: duplication.

**Table S4: Sensitivity analysis of the effect of gene dosage measured by pLI on NVIQ in autistic probands from SSC and MSSNG, and in the unselected population**.

| **Phenotype** | **Models** | **N** | **pLI DEL** | | | **pLI DUP** | | | |
| --- | --- | --- | --- | --- | --- | --- | --- | --- | --- |
|  |  |  | β | SE | p | | β | SE | p |
| NVIQ | All individuals | 6,656 | **-0.18** | **0.02** | **1.44×10^-16^** | | **-0.04** | **0.01** | **3.20×10^-3^** |
|  |  |  | Interaction pLI deletions and diagnosis p=0.86 | | | | Interaction pLI duplications and diagnosis p=0.16 | | |
|  |  |  | Interaction pLI deletions and sex p=0.62 | | | | Interaction pLI duplications and sex p=0.51 | | |
|  | Remove carriers of CNVs  with a pLI > 10 | 6,629 | **-0.18** | **0.03** | **2.43×10^-11^** | | -0.02 | 0.02 | 0.31 |
|  | Remove carriers of recurrent CNVs previously associated with NDD | 6,484 | **-0.20** | **0.03** | **3.42×10^-11^** | | **-0.05** | 0.02 | **9.53×10^-3^** |
|  | Remove carriers of rare *denovo* CNVs^(a)^ | 4,126 | **-0.11** | **0.05** | **0.04** | | -0.04 | 0.02 | 0.23 |

^(a)^ Information about the transmission of each CNV is not available in IMAGEN, the selection of rare CNVs are described in supplementary Methods. SE: Standard error; NVIQ: Non-verbal intelligence quotient; NDD: Neurodevelopmental disorders; pLI: probability of being Loss-of-function Intolerant; pLI DEL or pLI DUP: deleted or duplicated point of pLI score; CNV: Copy number variants.

**Table S5: Sensitivity analysis of the effect of gene dosage measured by pLI on autism risk using autistic probands from SSC and MSSNG, unaffected siblings from SSC, and the unselected population**.

| **Populations** | **Models** | **N individuals** | | **CNV score** | **Not adjusted**  **for NVIQ** | | | **Adjusted**  **for NVIQ** | | | |
| --- | --- | --- | --- | --- | --- | --- | --- | --- | --- | --- | --- |
|  |  | Probands | Controls |  | OR | 95%CI | p | | OR | 95%CI | p |
| **Probands (SSC-MSSNG)**  **Vs.**  **Unselected population** | All individuals | 3,703 | 2,224 | pLI DEL | **1.43** | **1.26-1.67** | **6.38×10^-7^** | | **1.28** | **1.11-1.53** | **2.14×10^-3^** |
|  |  |  |  | pLI DUP | **1.23** | **1.14-1.34** | **7.19×10^-7^** | | **1.21** | **1.11-1.34** | **3.56×10^-5^** |
|  |  |  |  | pLI DEL*sex | 1.01 | 0.76-1.37 | 0.96 | | 1.02 | 0.74-1.44 | 0.89 |
|  |  |  |  | pLI DUP*sex | 1.06 | 0.90-1.26 | 0.51 | | 1.07 | 0.89-1.29 | 0.50 |
|  | Remove carriers of CNVs with a pLI >10 | 3,680 | 2,222 | pLI DEL | **1.42** | **1.24-1.66** | **2.05×10^-6^** | | **1.28** | **1.10-1.52** | **2.95×10^-3^** |
|  |  |  |  | pLI DUP | **1.26** | **1.15-1.39** | **3.01×10^-6^** | | **1.24** | **1.12-1.39** | **8.43×10^-5^** |
|  | Remove carriers of recurrent CNVs associated with NDD | 3,578 | 2,193 | pLI DEL | **1.40** | **1.19-1.73** | **4.19×10^-4^** | | 1.21 | 0.99-1.56 | 0.10 |
|  |  |  |  | pLI DUP | **1.23** | **1.11-1.38** | **2.41×10^-4^** | | **1.21** | **1.08-1.38** | **2.09×10^-3^** |
|  | Remove carriers of rare *de-novo* CNVs^(a,b)^ | 3,070 | 480 | pLI DEL | 1.22 | 0.91-1.84 | 0.26 | | 1.20 | 0.86-1.90 | 0.34 |
|  |  |  |  | pLI DUP | 0.92 | 0.82-1.05 | 0.22 | | 0.89 | 0.78-1.02 | 0.08 |
|  | Removing the 10 individuals with autism risk from Imagen^(c)^ | 3,703 | 2,091 | pLI DEL | **1.41** | **1.24-1.64** | **1.49×10^-6^** | | **1.26** | **1.09-1.50** | **3.46×10^-3^** |
|  |  |  |  | pLI DUP | **1.22** | **1.13-1.33** | **1.98×10^-6^** | | **1.20** | **1.10-1.33** | **7.99×10^-5^** |
| **SSC probands**  **Vs. Siblings** | Remove carriers of rare *de-novo* CNVs^(b,d)^ | 1,950 | 1,950 | pLI DEL | **1.44** | **1.03-2.01** | **0.03** | | N.A. | N.A. | N.A. |
|  |  |  |  | pLI DUP | **1.21** | **1.03-1.41** | **0.02** | | N.A. | N.A. | N.A. |
|  |  | Unaffected siblings | Imagen + SYS |  |  |  |  | |  |  |  |
| **SSC unaffected siblings vs. Unselected population** | All individuals | 2,074 | 2,224 | pLI DEL | 1.03 | 0.88-1.21 | 0.69 | | N.A. | N.A. | N.A. |
|  |  |  |  | pLI DUP | 1.09 | 0.98-1.21 | 0.10 | | N.A. | N.A. | N.A. |

^(a)^ Information about the transmission of each CNV was not available in IMAGEN, this analysis was underpowered because of the lack of information on *de-novo* CNVs in the unselected population; ^(b)^ the selection of rare CNVs are described in supplementary Methods, ^(c)^ 10 individuals from IMAGEN met criteria for Autism as estimated by the DAWBA (Development and Well-Being Assessment), we also excluded 124 individual without diagnostic information (DAWBA), ^(d)^ NVIQ was not available in unaffected siblings for the adjustment. OR: Odds ratio; 95%CI: 95% Confidence interval; CNV: Copy number variant; NVIQ: Non-verbal IQ; NDD: Neurodevelopmental disorders; pLI: probability of being Loss-of-function Intolerant; pLI DEL or pLI DUP: deleted or duplicated point of pLI score; pLI DEL*sex or DUP*sex: interaction between pLI and sex; N.A.: Not applicable.

**Table S6: Breakpoints used to detect recurrent CNVs associated to autism and corresponding empirical and estimated odds ratios in autistic population.**

| **Locus** | **Type** | **Chr** | **Start - Stop hg19 (Mb)** | **Sum of pLI** | **N autism cases^(a)^** | **N**  **Controls**  **^(a)^** | **Published**  **autism risk ^(a)^** | | **Ref** | **Estimated autism risk^(b)^** | | **Estimated loss of NVIQ points** | | **Estimated gain of SRS points** | |
| --- | --- | --- | --- | --- | --- | --- | --- | --- | --- | --- | --- | --- | --- | --- | --- |
|  |  |  |  |  |  |  | OR | 95%CI |  | OR | 95%CI |  | 95%CI |  | 95%CI |
| 1q21.1  (class I) | DEL | 1 | 146.57-147.50 | 2.49 | 1/3,032 | 16/75,505 | **1.56** | **0.21-11.74** | (66) | **2.15** | **1.57-2.93** | 6.72 | 6.13-7.31 | 9.27 | 8.16-10.38 |
| 1q21.1  (class I) | DUP | 1 | 146.57-147.50 | 2.49 | 8/3,032 | 19/57,730 | 8.03 | 3.51-18.37 | (66) | 1.44 | 1.19-1.74 | 1.49 | 1.20-1.78 | 4.65 | 3.81-5.49 |
| 3q29 | DEL | 3 | 195.73-197.34 | 6.56 | 1/2,120 | 1/63,649 | **30.04** | **1.88-480.40** | (66) | **7.50** | **3.31-17.01** | 17.71 | 17.12-18.30 | 24.42 | 23.31-25.53 |
| 5q35 | DEL | 5 | 175.65-176.99 | 11.92 | 1/3,955 | 0/13,696 | ∞ | N.S. | (64) | 38.91 | 8.79-172.24 | 32.18 | 31.59-32.77 | 44.38 | 43.27-45.49 |
| 7q11.23  (WBS) | DUP | 7 | 72.72-74.15 | 10.27 | 4/2,120 | 1/16,257 | **30.73** | **3.43-275.07** | (66) | **4.45** | **2.03-9.72** | 6.16 | 5.87-6.45 | 19.16 | 18.32-20.00 |
| 15q11.2  (BP1-BP2) | DEL | 15 | 22.75-23.27 | 1.70 | 8/2,525 | 19/7,086 | **1.30** | **0.42-3.96** | (67) | **1.69** | **1.36-2.08** | 4.59 | 4.00-5.18 | 6.33 | 5.22-7.44 |
| 15q11.2  (BP1-BP2) | DUP | 15 | 22.75-23.27 | 1.70 | 20/2,525 | 38/7,086 | **1.80** | **0.82-3.97** | (67) | **1.28** | **1.12-1.46** | 1.02 | 0.73-1.31 | 3.17 | 2.33-4.01 |
| 15q13.3  (BP4-BP5) | DEL | 15 | 30.92-32.51 | 1.71 | 4/2,120 | 13/74,106 | 10.77 | 3.51-33.07 | (66) | 1.69 | 1.37-2.09 | 4.62 | 4.03-5.21 | 6.37 | 5.26-7.48 |
| 15q13.3  (BP4-BP5) | DUP | 15 | 30.92-32.51 | 1.71 | 2/3,955 | 5/13,696 | **1.39** | **0.27-7.14** | (64) | **1.28** | **1.13-1.46** | 1.03 | 0.74-1.32 | 3.19 | 2.35-4.03 |
| 16p11.2  (BP4-BP5) | DEL | 16 | 29.60-30.30 | 9.92 | 18/4,315 | 25/56,752 | **9.50** | **5.18-17.43** | (66) | **21.05** | **6.10-72.6** | 26.78 | 26.19-27.37 | 36.93 | 35.82-38.04 |
| 16p11.2  (BP4-BP5) | DUP | 16 | 29.60-30.30 | 9.92 | 17/4,315 | 19/56,752 | **11.81** | **6.13-22.74** | (66) | **4.23** | **1.99-9.00** | 5.95 | 5.66-6.24 | 18.51 | 17.67-19.35 |
| 16p11.2  distal | DEL | 16 | 28.81-29.04 | 3.82 | 1/3,955 | 2/13,696 | **1.73** | **0.16-19.10** | (64) | **3.23** | **2.01-5.21** | 10.31 | 9.72-10.90 | 14.22 | 13.11-15.33 |
| 16p11.2  distal | DUP | 16 | 28.81-29.04 | 3.82 | 1/3,955 | 3/13,696 | **1.15** | **0.12-11.10** | (64) | **1.74** | **1.30-2.33** | 2.29 | 2.00-2.58 | 7.13 | 6.29-7.97 |
| 16p13.11 | DEL | 16 | 15.12-16.29 | 2.84 | 5/3,955 | 4/13,696 | **4.34** | **1.16-16.14** | (64) | **2.39** | **1.68-3.41** | 7.67 | 7.08-8.26 | 10.57 | 9.46-11.68 |
| 16p13.11 | DUP | 16 | 15.12-16.29 | 2.84 | 4/2,120 | 81/62,973 | **1.47** | **0.54-4.01** | (66) | **1.51** | **1.22-1.88** | 1.70 | 1.41-1.99 | 5.30 | 4.46-6.14 |
| 17p11.2  (SMS) | DEL | 17 | 16.58-20.33 | 12.21 | 2/4,687 | 0/151,619 | ∞ | N.S. | (68) | 42.54 | 9.27-195.22 | 32.97 | 32.38-33.56 | 45.46 | 44.35-46.57 |
| 17p11.2  (SMS) | DUP | 17 | 16.58-20.33 | 12.21 | 1/4,687 | 1/151,619 | **32.30** | **2.02-517.39** | (68) | **5.90** | **2.33-14.94** | 7.33 | 7.04-7.62 | 22.79 | 21.95-23.63 |
| 17p12 | DEL | 17 | 14.04-15.41 | 1.17 | 2/2,120 | 14/59,086 | **4.00** | **0.9-17.5** | (66) | **1.43** | **1.24-1.66** | 3.16 | 2.57-3.75 | 4.36 | 3.25-5.47 |
| 17q12 | DEL | 17 | 34.81-36.25 | 5.07 | 2/2,120 | 4/68,131 | **16.08** | **2.94-87.86** | (66) | **4.75** | **2.52-8.94** | 13.69 | 13.10-14.28 | 18.87 | 17.76-19.98 |
| 22q11.2 | DEL | 22 | 18.89-21.90 | 11.44 | 5/4,687 | 5/151,619 | **32.37** | **9.37-111.87** | (68) | **33.58** | **8.05-139.99** | 30.89 | 30.30-31.48 | 42.59 | 41.48-43.70 |
| 22q11.2 | DUP | 22 | 18.89-21.90 | 11.44 | 12/4,315 | 23/27,133 | **3.28** | **1.63-6.61** | (66) | **5.27** | **2.21-12.60** | 6.86 | 6.57-7.15 | 21.35 | 20.51-22.19 |

^(a)^Based on number of carriers in autistic probands and controls reported in Malhotra et al. (2012) (66) Moreno DeLuca et al. (2013) (64), Chaste et al. (2014) (67), and Sanders et al. (2019) (68). In bold: CNVs for which previously published data and our estimation are overlapping. BP: Break points; WBS: William Beuren Syndrome; SMS: Smith Magenis Syndrome; DEL: deletion; DUP: duplication; chr: chromosome; hg19: Homo sapiens (human) genome assembly GRCh37 (hg19) from Genome Reference Consortium; pLI: probability of being Loss-of-function Intolerant; Sum of pLI: sum of score of pLI for the corresponding region; OR: Odds ratio; 95% CI: 95% confidence intervals; N.S.: Non-significant; Ref: reference.

**Table S7: Description of models used for the investigation of the effect of gene dosage on phenotypical measures of autistic probands from SSC**.

| **Phenotype** | **N** | **Type of normalization** | **Regression model** | **covariates** |
| --- | --- | --- | --- | --- |
| Autism related symptoms | | | | |
| **Regression** | 2,568 | N.A. | Logistic | Sex, ancestry |
| Language and phonology | | | | |
| **CTOPP** | 1,988 | z-scored with normative data: mean=10, SD=3 | Linear | Sex, ancestry |
| **Word delay** | 2,567 | N.A. | Logistic | Sex, ancestry |
| **Phrase delay** | 2,567 | N.A. | Logistic | Sex, ancestry |
| Adaptive skills (VABS-II) | | | | |
| **Total score** | 2,569 | z-scored with normative data: mean=100, SD=15 | Linear | Ancestry |
| **Daily living** | 2,569 | z-scored with normative data: mean=100, SD=15 | Linear | Ancestry |
| **Communication** | 2,569 | z-scored with normative data: mean=100, SD=15 | Linear | Ancestry |
| **Socialization** | 2,569 | z-scored with normative data: mean=100, SD=15 | Linear | Ancestry |
| Motor skills | | | | |
| **Motor VABS-II** | 919 | z-scored with normative data: mean=100, SD=15 | Linear | Ancestry |
| **Gross motor VABS-II** | 926 | z-scored with normative data: mean=15, SD=3 | Linear | Ancestry |
| **Fine motor**  **VABS-II** | 923 | z-scored with normative data: mean=15, SD=3 | Linear | Ancestry |
| **Delayed onset for walking** | 2,564 | N.A. | Logistic | Sex, ancestry |
| **Age of onset for walking** | 2,564 | N.A. | Quasi-Poisson | Sex, ancestry |
| **DCDQ score** | 2,209 | z-scored with probands data: mean=38.5, SD=12.4 | Linear | Age, sex, ancestry |
| Associated neurological condition | | | | |
| **Non-febrile seizure** | 2,566 | N.A. | Logistic | Sex, ancestry |

SD: Standard deviation; N.A.: Not applicable; CTOPP: Comprehensive Test of Phonological Processing; VABS-II: Vineland Adaptive behaviour Rating Scales - Second Edition; DCDQ: Developmental Coordination Disorder Questionnaire.

**Table S8**: **Effect of gene dosage measured by pLI on SRS using autistic probands from SSC, their unaffected siblings and parents and the unselected population from IMAGEN**.

| **Population** | **N** | **SRS-score** | **Model** | **CNV score** | **Effect size**  **(β or OR)** | **SE or 95%CI** | **p** |
| --- | --- | --- | --- | --- | --- | --- | --- |
| SSC probands | 2,556 | Total-raw | Linear not adjusted for NVIQ | pLI DEL | -0.21 | 0.40 | 0.60 |
|  |  |  |  | pLI DUP | -0.28 | 0.36 | 0.42 |
|  |  |  | Linear adjusted for NVIQ | pLI DEL | -0.31 | 0.41 | 0.45 |
|  |  |  |  | pLI DUP | -0.32 | 0.36 | 0.36 |
| SSC unaffected siblings | 2,078 | √Total-raw^(a)^ | Linear not adjusted for NVIQ | pLI DEL | 0.05 | 0.08 | 0.47 |
|  |  |  |  | pLI DUP | 0.001 | 0.06 | 0.99 |
| SSC parents | 4,838 | √Total-raw^(a)^ | Linear not adjusted for NVIQ | pLI DEL | 0.07 | 0.09 | 0.43 |
|  |  |  |  | pLI DUP | 0.01 | 0.04 | 0.83 |
| MSSNG probands | 598 | Total-raw | Linear not adjusted for NVIQ | pLI DEL | 0.71 | 1.78 | 0.69 |
|  |  |  |  | pLI DUP | -0.33 | 1.37 | 0.81 |
|  |  |  | Linear adjusted for NVIQ | pLI DEL | -0.70 | 1.76 | 0.69 |
|  |  |  |  | pLI DUP | -0.45 | 1.34 | 0.73 |
|  |  |  |  | NVIQ | **-0.30** | **0.06** | **1.32x10^-7^** |
| IMAGEN | 977 | √Total-raw^(a)^ | Linear not adjusted for NVIQ | pLI DEL | -0.06 | 0.15 | 0.66 |
|  |  |  |  | pLI DUP | 0.03 | 0.09 | 0.71 |
|  |  |  | Linear adjusted for NVIQ | pLI DEL | -0.09 | 0.15 | 0.56 |
|  |  |  |  | pLI DUP | 0.02 | 0.09 | 0.81 |
| SSC probands +  MSSNG probands | 3,154 | Total-raw | Linear not adjusted for NVIQ | pLI DEL | 0.44 | 0.51 | 0.39 |
|  |  |  |  | pLI DUP | -0.27 | 0.45 | 0.54 |
|  |  |  | Linear adjusted for NVIQ | pLI DEL | -0.41 | 0.50 | 0.41 |
|  |  |  |  | pLI DUP | -0.52 | 0.43 | 0.23 |
|  |  |  |  | NVIQ | **-0.27** | **0.02** | **1.40x10^-46^** |
| SSC probands +  MSSNG probands + IMAGEN | 4,131 | Total-raw | Linear not adjusted for NVIQ or autism diagnosis | pLI DEL | **3.66** | **0.56** | **5.53x10^-11^** |
|  |  |  |  | pLI DUP | **1.64** | **0.42** | **1.12x10^-3^** |
|  |  |  | Linear adjusted for autism diagnosis | pLI DEL | 0.56 | 0.35 | 0.11 |
|  |  |  |  | pLI DUP | -0.12 | 0.27 | 0.66 |
|  |  |  | Linear adjusted for NVIQ | pLI DEL | 0.42 | 0.59 | 0.46 |
|  |  |  |  | pLI DUP | 0.17 | 0.49 | 0.75 |
|  |  |  |  | NVIQ | **-0.52** | **0.02** | **1.26x10^-135^** |
|  |  |  | Linear adjusted for autism diagnosis and NVIQ | pLI DEL | -0.35 | 0.46 | 0.45 |
|  |  |  |  | pLI DUP | -0.45 | 0.39 | 0.25 |
|  |  |  |  | NVIQ | **-0.26** | **0.02** | **2.63x10^-52^** |
| Unaffected siblings + Unaffected Parents + IMAGEN | 7,926 | Total-raw | Linear mixed-effect | pLI DEL | 0.62 | 0.62 | 0.32 |
|  |  |  |  | pLI DUP | 0.08 | 0.36 | 0.82 |
| SSC probands + Unaffected siblings +  Unaffected Parents | 9,473 | Total-raw | Linear mixed-effect not adjusted for autism diagnosis | pLI DEL | **3.47** | **0.58** | **2.40x10^-9^** |
|  |  |  |  | pLI DUP | **1.54** | **0.44** | **5.20x10^-4^** |
|  |  |  | Linear mixed-effect adjusted for autism diagnosis | pLI DEL | 0.75 | 0.37 | 4.30x10^-2^ |
|  |  |  |  | pLI DUP | -0.003 | 0.29 | 0.99 |
| All SSC + IMAGEN | 10,483 | Total-raw | Linear mixed-effect not adjusted for autism diagnosis | pLI DEL | **3.72** | **0.57** | **5.10x10^-11^** |
|  |  |  |  | pLI DUP | **1.87** | **0.43** | **1.40x10^-5^** |
|  |  |  | Linear mixed-effect adjusted for autism diagnosis | pLI DEL | 0.55 | 0.36 | 0.13 |
|  |  |  |  | pLI DUP | -0.10 | 0.27 | 0.72 |
| All SSC +  MSSNG + IMAGEN | 11,081 | Total-raw | Linear mixed-effect not adjusted for autism diagnosis | pLI DEL | **3.68** | **0.56** | **4.30x10^-11^** |
|  |  |  |  | pLI DUP | **1.63** | **0.42** | **1.20x10^-4^** |
|  |  |  | Linear mixed-effect adjusted for autism diagnosis | pLI DEL | 0.56 | 0.35 | 0.11 |
|  |  |  |  | pLI DUP | -0.12 | 0.27 | 0.66 |
| Probands + Unaffected siblings + IMAGEN | 5,189 | SRS categories (normal, clinical) ^(b)^ | Logistic regression not adjusted for autism diagnosis | pLI DEL | **1.20** | **1.10-1.33** | **8.46x10^-^⁵** |
|  |  |  |  | pLI DUP | **1.13** | **1.06-1.22** | **2.00x10^-^⁴** |
|  |  |  | Logistic regression adjusted for autism diagnosis | pLI DEL | 0.96 | 0.84- 1.15 | 0.59 |
|  |  |  |  | pLI DUP | 0.97 | 0.86-1.12 | 0.66 |
| Probands + Unaffected siblings + IMAGEN | 5,189 | SRS categories (normal, moderate, mild, clinically significant) ^(b)^ | Ordinal cumulative not adjusted for autism diagnosis | pLI DEL | **1.20** | **1.11- 1.30** | **3.92x10^-^⁶** |
|  |  |  |  | pLI DUP | **1.11** | **1.05-1.18** | **3.00x10^-^⁴** |
|  |  |  | Ordinal cumulative adjusted for autism diagnosis | pLI DEL | 1.05 | 0.96-1.15 | 0.26 |
|  |  |  |  | pLI DUP | 0.99 | 0.93- 1.06 | 0.87 |

All linear, logistic, or ordinal regression models used were adjusted for age, sex and ancestry. Models take into account family as random-effect when including related individuals (Methods). Effect size are presented as β for linear regression models and as odds ratio for logistic and ordinal regression models. ^(a)^Square root transformation of the total SRS raw score was performed to adjust for the non-gaussian distribution or bimodality of SRS distribution (Figure 4); ^(b)^ Based on the previously published *T*-score categorization (55) (Methods). The statistical threshold after correction for multiple testing is p ≤ 2.7.10^-3^. Significant results are in bold. SE: Standard error; OR: Odds ratio; 95%CI: 95% Confidence interval; NVIQ: Non-verbal intelligence quotient; DEL: deletion; DUP: duplication; pLI: probability of being Loss-of-function Intolerant; pLI DEL or pLI DUP: deleted or duplicated point of pLI score; √Total-raw: square root transformation of the total SRS raw.

**Table S9: Effect of gene dosage measured by pLI on autism severity scores (main domains of ADI-R and ADOS-calibrated severity scores) using autistic probands from SSC and MSSNG.**

| **Phenotype** | **N** | **CNV score** | **Not adjusted for NVIQ** | | | **Adjusted for NVIQ** | | |
| --- | --- | --- | --- | --- | --- | --- | --- | --- |
|  |  |  | OR | 95%CI | p | OR | 95%CI | p |
| Probands from SSC | | | | | | | | |
| ADI-R  reciprocal social interactions | 2,567 | pLI DEL | 1.00 | 0.94-1.07 | 0.89 | 0.93 | 0.87-0.99 | 0.04 |
|  |  | pLI DUP | 1.06 | 1.01-1.11 | 0.02 | 1.03 | 0.98-1.08 | 0.20 |
| ADI-R  rrsb | 2,567 | pLI DEL | 0.95 | 0.89-1.02 | 0.14 | 0.94 | 0.88-1.01 | 0.10 |
|  |  | pLI DUP | 1.01 | 0.97-1.06 | 0.54 | 1.01 | 0.97-1.06 | 0.61 |
| ADI-R  verbal communication | 2,254 | pLI DEL | 1.00 | 0.93-1.08 | 0.91 | 0.95 | 0.88-1.02 | 0.18 |
|  |  | pLI DUP | 1.06 | 1.00-1.12 | 0.05 | 1.04 | 0.98-1.10 | 0.17 |
| ADI-R  non-verbal communication | 2,567 | pLI DEL | 1.01 | 0.95-1.08 | 0.73 | 0.94 | 0.88-1.00 | 0.07 |
|  |  | pLI DUP | 1.04 | 0.99-1.09 | 0.09 | 1.01 | 0.97-1.06 | 0.55 |
| ADOS  overall css^(a)^ | 2,499 | pLI DEL | 0.94 | 0.88-1.01 | 0.09 | 0.92 | 0.86-0.98 | 0.02 |
|  |  | pLI DUP | 1.02 | 0.97-1.07 | 0.41 | 1.01 | 0.96-1.06 | 0.73 |
| ADOS  social affect css^(a)^ | 2,371 | pLI DEL | 0.96 | 0.92-1.03 | 0.25 | 0.94 | 0.87-1.01 | 0.08 |
|  |  | pLI DUP | 1.04 | 0.99-1.10 | 0.08 | 1.03 | 0.98-1.08 | 0.19 |
| ADOS  rrsb css^(a)^ | 2,450 | pLI DEL | 0.98 | 0.91-1.05 | 0.55 | 0.95 | 0.88-1.02 | 0.12 |
|  |  | pLI DUP | 1.01 | 0.96-1.06 | 0.67 | 0.99 | 0.98-1.10 | 0.85 |
| ADI-R  Overall level of language | 2,568 | pLI DEL | 1.08 | 0.98-1.19 | 0.13 | 0.93 | 0.83-1.04 | 0.18 |
|  |  | pLI DUP | 1.06 | 1.00-1.13 | 0.04 | 1.03 | 0.97-1.10 | 0.36 |
| Probands from MSSNG | | | | | | | | |
| ADI-R  reciprocal social interactions | 397 | pLI DEL | 1.26 | 0.85-1.85 | 0.25 | 1.20 | 0.82-1.77 | 0.35 |
|  |  | pLI DUP | 0.98 | 0.77-1.25 | 0.85 | 0.99 | 0.77-1.26 | 0.93 |
| ADI-R  rrsb | 695 | pLI DEL | 1.15 | 0.98-1.37 | 0.09 | 1.15 | 0.97-1.36 | 0.11 |
|  |  | pLI DUP | 1.00 | 0.86-1.13 | 0.87 | 0.99 | 0.86-1.13 | 0.83 |
| ADI-R  verbal communication | 370 | pLI DEL | 1.08 | 0.82-1.42 | 0.59 | 1.04 | 0.79-1.37 | 0.77 |
|  |  | pLI DUP | 1.01 | 0.79-1.29 | 0.95 | 0.97 | 0.76-1.24 | 0.80 |
| ADI-R  non-verbal communication | 73 | pLI DEL | 1.12 | 0.77-1.64 | 0.49 | 1.10 | 0.75-1.61 | 0.63 |
|  |  | pLI DUP | 0.90 | 0.75-1.07 | 0.27 | 0.89 | 0.74-1.07 | 0.21 |
| ADOS  overall css^(a)^ | 733 | pLI DEL | 0.94 | 0.75-1.18 | 0.58 | 0.92 | 0.73-1.16 | 0.50 |
|  |  | pLI DUP | 0.86 | 0.75-0.99 | 0.04 | 0.86 | 0.75-0.99 | 0.04 |
| ADOS  social affect css^(a)^ | 373 | pLI DEL | 0.74 | 0.50-1.11 | 0.15 | 0.74 | 0.50-1.11 | 0.15 |
|  |  | pLI DUP | 0.90 | 0.76-1.07 | 0.23 | 0.90 | 0.76-1.07 | 0.23 |
| ADOS  rrsb css^(a)^ | 388 | pLI DEL | 0.91 | 0.61-1.36 | 0.64 | 0.91 | 0.61-1.36 | 0.63 |
|  |  | pLI DUP | 0.95 | 0.83-1.12 | 0.60 | 0.96 | 0.82-1.11 | 0.57 |
| ADI-R  Overall level of language | 1,267 | pLI DEL | 1.27 | 1.03-1.21 | 7.19.10^-3^ | 1.13 | 0.94-1.37 | 0.19 |
|  |  | pLI DUP | 1.05 | 1.03-1.21 | 0.49 | 1.05 | 0.92-1.20 | 0.44 |
| Pooled probands (SSC + MSSNG) | | | | | | | | |
| ADI-R  reciprocal social interactions | 2,966 | pLI DEL | 1.01 | 0.95-1.08 | 0.69 | 0.94 | 0.88-1.01 | 0.08 |
|  |  | pLI DUP | 1.05 | 1.00-1.10 | 0.03 | 1.03 | 0.98-1.08 | 0.24 |
| ADI-R  rrsb | 3,264 | pLI DEL | 0.98 | 0.92-1.04 | 0.52 | 0.97 | 0.91-1.04 | 0.40 |
|  |  | pLI DUP | 1.01 | 0.97-1.06 | 0.58 | 1.01 | 0.97-1.06 | 0.66 |
| ADI-R  verbal communication | 2,626 | pLI DEL | 1.01 | 0.94-1.08 | 0.80 | 0.95 | 0.89-1.03 | 0.20 |
|  |  | pLI DUP | 1.06 | 0.99-1.12 | 0.05 | 1.04 | 0.98-1.10 | 0.20 |
| ADI-R  non-verbal communication | 2,642 | pLI DEL | 1.02 | 0.95-1.09 | 0.65 | 0.94 | 0.88-1.01 | 0.09 |
|  |  | pLI DUP | 1.03 | 0.99-1.08 | 0.18 | 1.01 | 0.96-1.05 | 0.82 |
| ADOS  overall css^(a)^ | 3,122 | pLI DEL | 0.94 | 0.88-1.01 | 0.08 | 0.92 | 0.86-0.98 | 0.01 |
|  |  | pLI DUP | 0.99 | 0.95-1.04 | 0.88 | 0.99 | 0.94-1.03 | 0.56 |
| ADOS  social affect css^(a)^ | 2,675 | pLI DEL | 0.95 | 0.89-1.02 | 0.16 | 0.93 | 0.87-1.00 | 0.05 |
|  |  | pLI DUP | 1.03 | 0.98-1.08 | 0.25 | 1.02 | 0.97-1.07 | 0.44 |
| ADOS  rrsb css^(a)^ | 2,766 | pLI DEL | 0.98 | 0.91-1.05 | 0.50 | 0.94 | 0.88-1.01 | 0.11 |
|  |  | pLI DUP | 1.00 | 0.96-1.05 | 0.91 | 0.99 | 0.95-1.04 | 0.67 |
| ADI-R  Overall level of language | 3,607 | pLI DEL | 1.11 | 1.02-1.21 | 0.01 | 0.97 | 0.88-1.06 | 0.50 |
|  |  | pLI DUP | 1.06 | 1.00-1.12 | 0.03 | 1.04 | 0.98-1.10 | 0.23 |

All ordinal regression models used for each severity score were adjusted for age and sex, and ancestry when available (for autistic probands from SSC only). ^(a)^Calibrated severity score were computed based on previously published methodology from Hus et al. (2014) (40). The statistical threshold after correction for multiple testing is p ≤ 2.7.10^-3^. NVIQ: Non-verbal intelligence quotient; OR: Odds ratio; 95%CI: 95% Confidence interval; ADI-R: Autism Diagnostic Interview-Revised; ADOS: Autism Diagnostic Observation Schedule; css: calibrated severity score; rrsb: repetitive, restricted and stereotyped behaviours; DEL: deletion; DUP: duplication; pLI: probability of being Loss-of-function Intolerant; pLI DEL or pLI DUP: deleted or duplicated point of pLI score.

**Table S10**: **Effect of gene dosage measured by pLI on CBCL using autistic probands from SSC and unaffected siblings.**

| **Population** | **N** | **Model** | **CNV score** | **OR** | **95%CI** | **p** |
| --- | --- | --- | --- | --- | --- | --- |
| CBCL total Problems raw score | | | | | | |
| Probands | 1,945 | Negative Binomial not adjusted for NVIQ | pLI DEL | 1.01 | 0.99-1.03 | 0.43 |
|  |  |  | pLI DUP | 1.00 | 0.98-1.01 | 0.64 |
|  |  | Negative Binomial adjusted for NVIQ | pLI DEL | 1.01 | 0.99-1.03 | 0.40 |
|  |  |  | pLI DUP | 1.00 | 0.98-1.01 | 0.67 |
| Unaffected Siblings | 1,596 | Negative Binomial not adjusted for NVIQ | pLI DEL | 1.07 | 0.98-1.20 | 0.13 |
|  |  |  | pLI DUP | 1.05 | 0.99-1.13 | 0.10 |
| Probands + Unaffected siblings | 3,541 | Negative Binomial mixed-effect not adjusted for autism diagnosis | pLI DEL | **1.05** | **1.03-1.08** | **1.94x10^-6^** |
|  |  |  | pLI DUP | 1.02 | 1.01-1.04 | 3.04x10**^-3^** |
|  |  | Negative Binomial mixed-effect adjusted for autism diagnosis | pLI DEL | 1.01 | 1.00-1.03 | 0.11 |
|  |  |  | pLI DUP | 1.00 | 1.00-1.02 | 0.53 |
| CBCL externalizing Problems raw score | | | | | | |
| Probands | 1,945 | Negative Binomial not adjusted for NVIQ | pLI DEL | 1.01 | 0.98-1.05 | 0.37 |
|  |  |  | pLI DUP | 1.00 | 0.97-1.02 | 0.91 |
|  |  | Negative Binomial adjusted for NVIQ | pLI DEL | 1.01 | 0.98-1.05 | 0.41 |
|  |  |  | pLI DUP | 1.00 | 0.97-1.02 | 0.87 |
| Unaffected Siblings | 1,596 | Negative Binomial not adjusted for NVIQ | pLI DEL | 1.10 | 0.97-1.28 | 0.12 |
|  |  |  | pLI DUP | 1.04 | 0.95-1.13 | 0.40 |
| Probands + Unaffected siblings | 3,541 | Negative Binomial mixed-effect not adjusted for autism diagnosis | pLI DEL | **1.05** | **1.02-1.09** | **4.94x10^-4^** |
|  |  |  | pLI DUP | 1.02 | 1.00-1.05 | 0.02 |
|  |  | Negative Binomial mixed-effect adjusted for autism diagnosis | pLI DEL | 1.02 | 0.99-1.05 | 0.14 |
|  |  |  | pLI DUP | 1.01 | 0.98-1.03 | 0.56 |
| CBCL internalizing Problems raw score | | | | | | |
| Probands | 1,945 | Negative Binomial not adjusted for NVIQ | pLI DEL | 0.99 | 0.97-1.02 | 0.63 |
|  |  |  | pLI DUP | 0.98 | 0.96-1.00 | 0.07 |
|  |  | Negative Binomial adjusted for NVIQ | pLI DEL | 1.01 | 0.98-1.04 | 0.57 |
|  |  |  | pLI DUP | 0.99 | 0.96-1.01 | 0.19 |
| Unaffected Siblings | 1,596 | Negative Binomial not adjusted for NVIQ | pLI DEL | 1.08 | 0.97-1.22 | 0.15 |
|  |  |  | pLI DUP | 1.06 | 0.99-1.14 | 0.10 |
| Probands + Unaffected siblings | 3,541 | Negative Binomial mixed-effect not adjusted for autism diagnosis | pLI DEL | **1.04** | **1.01-1.07** | **2.12x10^-3^** |
|  |  |  | pLI DUP | 1.01 | 0.99-1.03 | 0.19 |
|  |  | Negative Binomial mixed-effect adjusted for autism diagnosis | pLI DEL | 1.01 | 0.98-1.03 | 0.61 |
|  |  |  | pLI DUP | 0.99 | 0.97-1.01 | 0.52 |

All negative binomial models used were adjusted for age, sex and ancestry. Models take into account family as random-effect when including related individuals (Methods). Effect size are presented as odds ratio. The statistical threshold after correction for multiple testing is p ≤ 2.7.10^-3^. Significant results are in bold. OR: Odds ratio; 95%CI: 95% Confidence interval; CBCL: Child behaviour Checklist; NVIQ: Non-verbal intelligence quotient; DEL: deletion; DUP: duplication; pLI: probability of being Loss-of-function Intolerant; pLI DEL or pLI DUP: deleted or duplicated point of pLI score.

**Table S11: Effect of gene dosage measured by pLI on general intelligence in autistic probands from SSC and MSSNG, and in the unselected population.**

| **Population** | **Measure** | **N** | **Covariates** | **CNV score** | **β** | **SE** | **p** |
| --- | --- | --- | --- | --- | --- | --- | --- |
| Probands (SSC) | NVIQ | 2,564 | Sex, type of test used, ancestry | pLI DEL | **-0.17** | **0.03** | **8.29×10^-10^** |
|  |  |  |  | pLI DUP | **-0.06** | **0.02** | **1.50×10^-3^** |
|  | NVIQ-DAS | 2,244 | Sex, ancestry | pLI DEL | **-0.16** | **0.03** | **1.20 ×10^-7^** |
|  |  |  |  | pLI DUP | **-0.07** | **0.02** | **1.10×10^-3^** |
|  | NVR | 1,958 | Sex, ancestry | pLI DEL | **-0.10** | **0.03** | **4.60×10^-4^** |
|  |  |  |  | pLI DUP | **-0.06** | **0.02** | **5.90×10^-3^** |
|  | Matrices | 1,958 | Sex, ancestry | pLI DEL | **-0.09** | **0.03** | **9.30 ×10^-4^** |
|  |  |  |  | pLI DUP | **-0.04** | **0.02** | **4.80×10^-2^** |
| Unselected | NVIQ | 2,710 | Sex, type of test used, ancestry | pLI DEL | **-0.19** | **0.04** | **6.90×10^-5^** |
|  |  |  |  | pLI DUP | 0.02 | 0.03 | 0.52 |
| Probands (MSSNG) | NVIQ | 1,381 | Sex, type of test used | **pLI DEL** | **-0.20** | **0.07** | **3.10×10^-3^** |
|  |  |  |  | pLI DUP | -0.02 | 0.04 | 0.61 |

All linear regression models were performed using a z-scored dependant variable. These z-scores are computed using normative data (e.g. mean NVIQ=100, SD NVIQ=15). CNV: Copy number variant; SE: Standard error; NVIQ: Non-verbal IQ; DAS: Differential Ability Scales; NVR: Non-verbal reasoning; pLI DEL or pLI DUP: deleted or duplicated point of pLI score.

**Table S12: Effect of gene dosage measured by pLI on autism risk.**

| **Group comparison** | **N** | | **pLI DEL** | | | | | **pLI DUP** | | | | |
| --- | --- | --- | --- | --- | --- | --- | --- | --- | --- | --- | --- | --- |
|  | probands | controls | Sum pLI in probands | Sum pLI in controls | OR | 95%CI | p | Sum pLI in probands | Sum pLI in controls | OR | 95%CI | p |
| Not adjusted for NVIQ | | | | | | | | | | | | |
| Probands vs. Unaffected Siblings | 2,074 | 2,074 | 346.58 | 117.67 | **1.43** | **1.23-1.66** | **3.78×10^-6^** | 640.32 | 371.62 | **1.32** | **1.17-1.49** | **4.63×10^-6^** |
| Probands vs. Unselected population | 2,569 | 2,223 | 416.00 | 114.11 | **1.40** | **1.23-1.64** | **2.33×10^-6^** | 816.99 | 330.31 | **1.30** | **1.19-1.42** | **1.88×10^-8^** |
| Adjusted for NVIQ | | | | | | | | | | | | |
| Probands vs. unselected population | 2,569 | 2,223 | 416.00 | 114.11 | **1.22** | **1.05-1.45** | **0.01** | 816.99 | 330.31 | **1.27** | **1.15-1.42** | **4.89×10^-6^** |
| Probands vs. unselected population | 1,438 | 1,438 | 153.80 | 93.51 | 1.21 | 1.01-1.45 | 60.4 %* | 357.01 | 219.40 | **1.25** | **1.09-1.43** | **98.2 %*** |
| Replication with probands from MSSNG | | | | | | | | | | | | |
| Not adjusted | | | | | | | | | | | | |
| Probands vs. Unselected population | 1,139 | 2,223 | 156.41 | 114.11 | **1.54** | **1.30-1.88** | **4.56×10^-6^** | 298.32 | 330.31 | **1.19** | **1.08-1.33** | **8.30×10^-4^** |
| Adjusted for NVIQ | | | | | | | | | | | | |
| Probands vs. unselected population | 1,139 | 2,223 | 156.41 | 114.11 | **1.39** | **1.15-1.72** | **1.44×10^-3^** | 298.32 | 330.31 | **1.20** | **1.08-1.35** | **1.44×10^-3^** |

Odds ratios are computed using a logistic regression including sum of pLI in deletions and duplications as the two main explanatory variables. OR represents the autism risk conferred for each deleted or duplicated point of pLI. Sum of pLI represents the sum of all genes deleted or duplicated in all individuals for the group. *percentage of matching iteration with a p-value ≤ 0.05 (for 500 iterations). Significative p-values are in bold (≤ 0.05). pLI: probability of being Loss-of-function Intolerant; sum of pLI: sum of score of pLI in the entire population; pLI DEL or pLI DUP: deleted or duplicated point of pLI; NVIQ: Non-verbal intelligence quotient; OR: Odds ratio; 95%CI: 95% Confidence interval.

**Table S13: Autism risk potentially mediated by NVIQ in the pooled dataset (SSC, MSSNG, unselected populations).**

| **Populations** | **N individuals** | | **Effect** | **Variable** | **OR** | **95% CI** | **p** | |
| --- | --- | --- | --- | --- | --- | --- | --- | --- |
|  | **probands** | **controls** |  |  |  |  |  |  |
| **Probands (SSC-MSSNG) *vs.* Unselected populations** | 3,703 | 2,224 | Direct effect | pLI DEL | **1.23** | **[1.08-1.41]** | | **2.28×10^-3^** |
|  |  |  |  | pLI DUP | **1.18** | **[1.10-1.26]** | | **1.00×10^-6^** |
|  |  |  | Indirect effect | pLI DEL | **1.17** | **[1.13-1.21]** | | **2.00×10^-16^** |
|  |  |  |  | pLI DUP | **1.06** | **[1.03-1.08]** | | **6.56×10^-7^** |
|  |  |  | Total effect | pLI DEL | **1.45** | **[1.26-1.65]** | | **7.33×10^-8^** |
|  |  |  |  | pLI DUP | **1.25** | **[1.16-1.34]** | | **2.92×10^-9^** |

Odds ratios are computed using a counterfactual-based mediation analysis on two different logistic regression including sum of pLI in deletions and duplications as the main explanatory variables. OR represents the autism risk conferred for each deleted or duplicated point of pLI. Direct effects of CNVs are those not mediated by NVIQ. Indirect are those potentially mediated by NVIQ. Total effects are those computed without adjusting for NVIQ. Significative p-values are in bold (≤ 0.05). pLI: probability of being Loss-of-function Intolerant. pLI DEL or pLI DUP: deleted or duplicated point of pLI. NVIQ: Non-verbal intelligence quotient; OR: Odds ratio; 95%CI: 95% Confidence interval.

**Table S14: Autism risk measured by pLI in subgroups of individual below or above the median (98) in the pooled dataset (SSC, MSSNG, unselected populations).**

| **Populations** | **N individuals** | | **IQ subgroup** | **Variable** | **OR** | **95% CI** | **p** | |
| --- | --- | --- | --- | --- | --- | --- | --- | --- |
|  | **probands** | **controls** |  |  |  |  |  |  |
| **Probands (SSC-MSSNG) *vs*. Unselected populations** | 2,363 | 667 | Below median NVIQ | pLI DEL | **1.27** | **[1.10-1.53]** | | **3.41×10^-3^** |
|  |  |  |  | pLI DUP | **1.34** | **[1.16-1.61]** | | **5.57×10^-4^** |
|  | 1,340 | 1,557 | Above median NVIQ | pLI DEL | **1.53** | **[1.19-2.07]** | | **2.17×10^-3^** |
|  |  |  |  | pLI DUP | **1.16** | **[1.04-1.31]** | | **1.10×10^-2^** |

Odds ratios are computed using a logistic regression including sum of pLI in deletions and duplications as the two main explanatory variables. OR represents the autism risk conferred for each deleted or duplicated point of pLI. Significative p-values are in bold (≤ 0.05). pLI: probability of being Loss-of-function Intolerant. pLI DEL or pLI DUP: deleted or duplicated point of pLI. NVIQ: Non-verbal intelligence quotient; OR: Odds ratio; 95%CI: 95% Confidence interval.

**Table S15: Estimated Genome wide effects of gene dosage on autism risk.**

|  | **ALL CNVs (including pLI=0)** | | | | **CNVs with pLI > 0** | | | |
| --- | --- | --- | --- | --- | --- | --- | --- | --- |
|  | >50Kb in autistic sample | | 1MB genome wide | | >50Kb in autistic sample | | 1MB genome wide | |
|  | OR DEL | OR DUP | OR DEL | OR DUP | OR DEL | OR DUP | OR DEL | OR DUP |
| N CNVs | 5,319 | 4,614 | 5,586 | 5,586 | 2,136 | 2,045 | 4,377 | 4,377 |
| Median | 1.00 | 1.00 | 1.36 | 1.21 | 1.02 | 1.05 | 1.58 | 1.32 |
| Mean | 1.15 | 1.13 | 3.05 | 1.58 | 1.36 | 1.31 | 3.62 | 1.75 |
| 70% | 1.00 | 1.00 | 1.85 | 1.45 | 1.02 | 1.17 | 2.13 | 1.58 |
| 75% | 1.01 | 1.02 | 2.04 | 1.54 | 1.09 | 1.2 | 2.38 | 1.69 |
| 80% | 1.02 | 1.05 | 2.35 | 1.68 | 1.09 | 1.21 | 2.66 | 1.82 |
| 90% | 1.09 | 1.21 | 3.72 | 2.23 | 1.35 | 1.33 | 4.43 | 2.48 |
| 95% | 1.31 | 1.33 | 6.65 | 3.17 | 1.60 | 1.63 | 8.28 | 3.63 |

pLI: probability of being Loss-of-function Intolerant; NVIQ: Non-verbal intelligence quotient; OR DEL/DUP: Estimated odds ratio for deletions or duplications based on pLI score. Odds ratio represents the estimated autism risk conferred by deleted or duplicated points of pLI in the autistic sample (probands from SSC and MSSNG) and in a genome of reference (Human Gene Nomenclature) (69) partitioned in CNV of 1Mb.
